# Arabidopsis lines with modified ascorbate concentrations reveal a link between ascorbate and auxin biosynthesis

**DOI:** 10.1101/2025.05.15.654287

**Authors:** M. Fenech, V. Zulian, J. Moya-Cuevas, D. Arnaud, I. Morilla, N. Smirnoff, M.A. Botella, A.N. Stepanova, J.M. Alonso, C. Martin-Pizarro, V. Amorim-Silva

**Affiliations:** Department of Plant and Microbial Biology, College of Agriculture and Life Sciences, North Carolina State University, 112 Derieux Place, Raleigh, North Carolina 27695-7614, United States of America; Department of Biological Sciences, Clemson University, 132 Long Hall, Clemson, South Carolina 29631, United States of America; Área de Mejora y Fisiología de Plantas, Instituto de Hortofruticultura Subtropical y Mediterránea “La Mayora”, Universidad de Málaga-Consejo Superior de Investigaciones Científicas (IHSM-UMA-CSIC), Universidad de Málaga, Campus Teatinos, 29010, Málaga, Spain; Foundation for Research and Technology Hellas, IMBB/FORTH, Institute of Molecular Biology and Biotechnology, N Plastira Str 100, Irakleio, 70013, Greece; Biosciences, Faculty of Life and Environmental Sciences, University of Exeter, Exeter, EX4 4QD, UK; Área de Protección de Cultivos, MLiMO, Instituto de Hortofruticultura Subtropical y Mediterránea “La Mayora”, Universidad de Málaga-Consejo Superior de Investigaciones Científicas (IHSM-UMA-CSIC), Universidad de Málaga, Campus Teatinos, 29010, Málaga, Spain; Université Sorbonne Paris Nord, LAGA, CNRS, UMR 7539, F-93430, Villetaneuse, France

**Author notes:** Corresponding authors. MF /, VAS.

## Abstract

Ascorbate is the most abundant water-soluble antioxidant in plants, and it is an essential molecule for normal plant development. It is present in all green plants, with very different concentrations in different plant species. While ascorbate accumulation is a trait of nutritional, and therefore, agronomical interest, the impact of different concentrations over cellular homeostasis remains elusive. In order to shed light over this question, we leveraged Arabidopsis lines with very low ascorbate (*vtc2* mutant with 20% of WT ascorbate levels), and low ascorbate concentration (*vtc4* mutant with 65% of WT levels), and we generated a line that accumulates 165% of WT levels (*vtc2/OE-VTC2*). An 80% reduction of ascorbate increased the expression of genes implicated in defense against pathogens, but repressed genes associated with abiotic stress responses. Unexpectedly, lines with increased (165% of WT) and decreased (65% of WT) ascorbate levels shared 85% of induced transcription factors and the GO terms associated with their transcriptional programs. Among the group of genes whose expression is positively correlated with ascorbate content, we identified *TAA1/WEI8*, a gene encoding a tryptophan aminotransferase that catalyzes the first step of auxin biosynthesis. Using a combination of genetic and pharmacological approaches in fluorescent and histochemical reporter lines for auxin biosynthesis and signaling activity, we revealed that TAA1- and TAA1-RELATED2 (TAR2)-mediated auxin biosynthesis is necessary for plants to cope with increased ascorbate concentration in a light-dependent manner, revealing a new layer of complexity in the regulatory landscape of redox homeostasis.

## Introduction

L-Ascorbate (ascorbate), also known as vitamin C, is the most abundant water-soluble antioxidant in plants. Humans, among other groups of animals, have lost the ability to synthesize ascorbate throughout evolution due to a lack of selective pressure as fruits and vegetables became a substantial part of our diet (Chatterjee, 1973; Smirnoff, 2018). Ascorbate deficiency in humans leads to scurvy, a disease considered rare although there are still numerous cases in modern society due to an increasing trend in choosing fast food (Lipner, 2018; Urueña-Palacio et al., 2018), with pediatric cases increasing by three-fold in the period 2016-2020 (Reikersdorfer et al., 2024). In addition to preventing scurvy, ascorbate is involved in many other processes in humans as a cofactor in various biological processes and it plays a major role in gene expression regulation (Blaschke et al., 2013; Minor et al., 2013; Hu et al., 2015; Young et al., 2015; Padayatty and Levine, 2016). The abovementioned reports suggest that in humans, ascorbate is not only critical for maintaining cell redox state but also for epigenetic regulation of gene expression and chromatin remodeling. Because the major source of vitamin C in humans is fresh produce, understanding how plants biosynthesize ascorbate, and how altered concentrations of this antioxidant impact plant development, is critical.

In plants, ascorbate is also a crucial molecule for normal development, with ascorbate-deprived mutants showing lethality that can be rescued by exogenous supplementation with ascorbate (Dowdle et al., 2007; Pineau et al., 2008; Lim et al., 2016; Fenech et al., 2021). Among its numerous roles described in plants, ascorbate is required to maintain the oxidative status in the active center of several enzymes like the Cu^2+^ and Fe^2+^/α-ketoglutarate-dependent dioxygenases—for example, 4-hydroxyphenylpyruvate dioxygenase involved in plastoquinone and tocopherol biosynthesis (Norris et al., 1998)—, modulating redox-sensitive processes such as anthocyanin biosynthesis (Page et al., 2012), as well as to scavenge reactive oxygen species (ROS) produced during photosynthesis (Tyystjärvi, 2008; Tóth et al., 2013), normal plant development (Wu et al., 2023), or in response to pathogens (Barth et al., 2004; Pavet et al., 2005; Mukherjee et al., 2010). Furthermore, ascorbate has been implicated in the crosstalk with other regulatory molecules, such as phytohormones, to help tune signaling cascades in response to various stresses (Akram et al., 2017; Veljović-Jovanović et al., 2017). Altered ascorbate status has been reported to influence salicylic acid, jasmonic acid, and abscisic acid (ABA) concentration (Pastori et al., 2003; Bulley et al., 2009). Interactions between indole-3-acetic acid (IAA) and ascorbate are suggested by ascorbate effects on root gravitropism (Lee et al., 2011) and root cell identity (Lee et al., 2007) and by a possible role for ascorbate oxidase in IAA decarboxylation and maintenance of the root quiescent center (Kerk et al., 2000; Jiang et al., 2003).

The major active form of auxin, IAA, is biosynthesized from the amino acid tryptophan (Trp) in two steps by two multi-gene families. First, Trp is converted into indole-3-pyruvic acid (IPyA) by a tryptophan aminotransferase encoded by three genes in *Arabidopsis thaliana* (hereon, Arabidopsis): *TRYPTOPHAN AMINOTRANSFERASE OF ARABIDOPSIS1 (TAA1)*, *TAA1-RELATED1* (*TAR1)*, and *TAR2* (Stepanova et al., 2008; Tao et al., 2008; Yamada et al., 2009). Then, a family of flavin monooxygenases, represented in Arabidopsis by *YUCCA1 (YUC1)* through *YUC11*, catalyzes the conversion of IPyA into IAA (Zhao et al., 2001; Stepanova et al., 2011). The spatiotemporal regulation of local auxin biosynthesis and the auxin transport patterns confer a very finely tuned control of auxin activity (Brumos et al., 2018; Brumos et al., 2020).

An interaction between auxin and ascorbate is supported not only by the abovementioned studies but also by the antagonistic effects of auxin (Mangano et al., 2017) and ascorbate (Tyburski et al., 2012) on ROS-mediated cell elongation of root hairs. While auxin induces *ROOT HAIR DEFECTIVE 6-LIKE4 (RSL4)*-mediated ROS production and root hair elongation (Mangano et al., 2017), high concentrations of ascorbate lead to shorter root hairs (Tyburski et al., 2012). Given the well-established role of ascorbate as a ROS scavenger (Foyer and Kunert, 2024), the antagonistic roles of IAA and ascorbate are thought to stem from their opposite effects on ROS production and neutralization (Wang et al., 2021). Accordingly, ascorbate biosynthesis must be finely tuned to ensure the right spatiotemporal distribution of ROS and normal plant development.

The rate-limiting step of ascorbate biosynthesis is catalyzed by GDP-L-galactose phosphorylase (GGP), which is encoded by *VITAMIN C DEFICIENT2* (*VTC2)* and *VTC5* in Arabidopsis (Fig. 1A) (Dowdle et al., 2007; Linster et al., 2007; Bulley et al., 2009; Fenech et al., 2021). The translation of *GGP* mRNA is controlled by an upstream open reading frame (uORF), which is proposed to mediate feedback repression by ascorbate (Laing et al., 2015; Baldet et al., 2024). *GGP* expression is controlled transcriptionally by light (Dowdle et al., 2007) and its activity is reversibly inhibited by a Per-ARNT-Sim/Light-Oxygen-Voltage (PAS/LOV) protein (PLP) photoreceptor when exposed to blue light in Arabidopsis (Aarabi et al., 2023) and tomato (Bournonville et al., 2023), thus linking light signaling and ascorbate production. In addition, ethylene and ABA regulate *VTC2* expression through ETHYLENE INSENSITIVE3 (EIN3) (positive regulator of *VTC2* expression) and ABA INSENSITIVE4 (ABI4) (negative regulator) to tune ascorbate concentration and, therefore, ROS levels (Yu et al., 2019). GGP is the first step in the pathway that does not produce cell wall precursors (Fig. 1A). Consistently, ascorbate biosynthesis mutants affected in steps upstream of GGP have cell wall and growth defects or even lethality at the gametophyte (Mounet-Gilbert et al., 2016; Qi et al., 2017) or embryo (Lukowitz et al., 2001) stages, while *ggp* and downstream mutants lethality can be prevented by exogenous supplementation of ascorbate (Dowdle et al., 2007; Pineau et al., 2008; Fenech et al., 2021).

**Figure 1.**
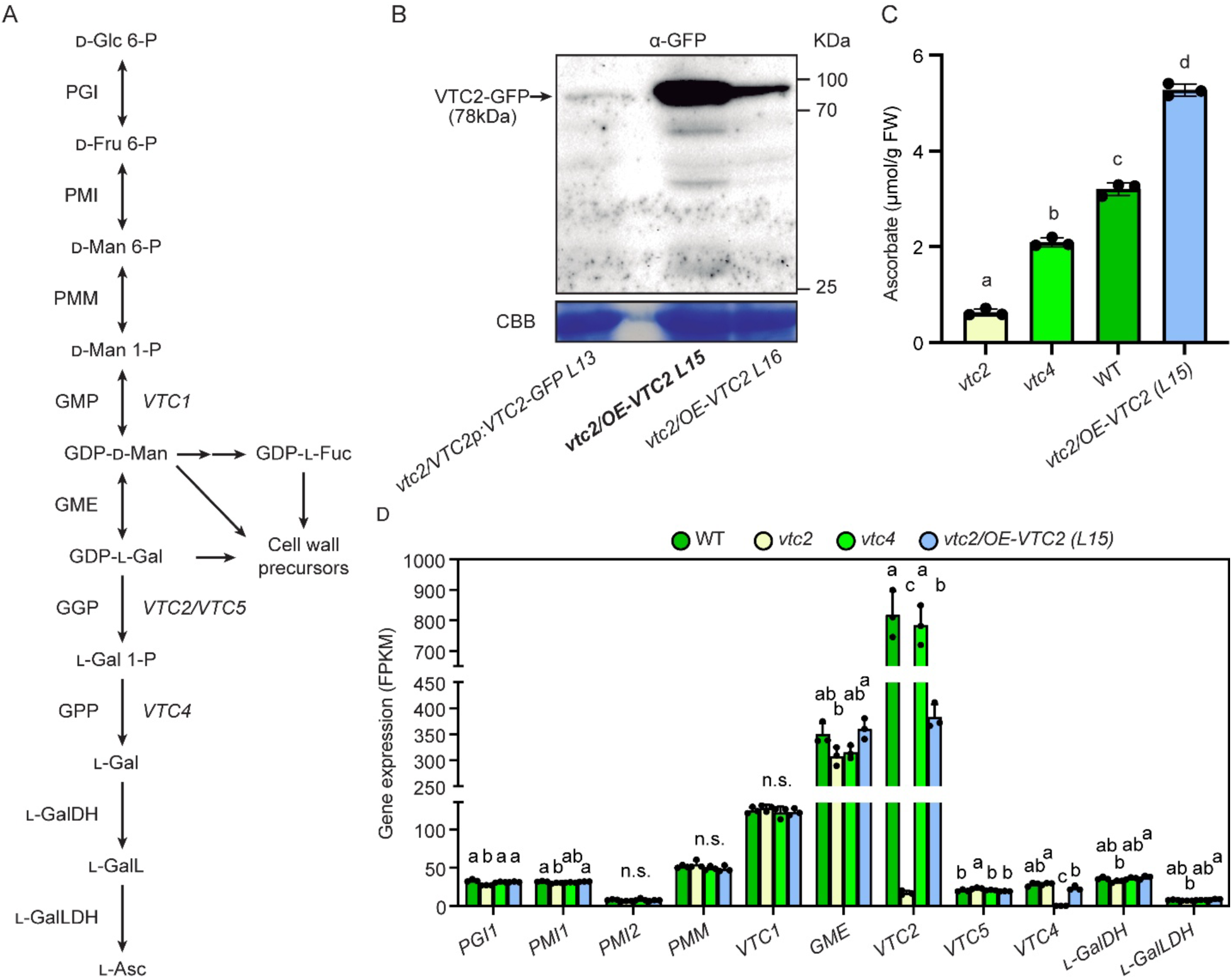
Arabidopsis ascorbate-altered lines do not show a compensatory response of ascorbate biosynthesis-related genes. A) Ascorbate biosynthesis (Smirnoff-Wheeler) pathway and its convergence with the biosynthesis of cell wall precursors. B) uORF-less *VTC2* overexpression lines accumulate more VTC2 protein than the native promoter line described in Fenech et al., 2021. C) A range of endogenous ascorbate concentrations (20% to 165% of WT levels) in four Arabidopsis lines employed in this work enables to study the effect of different ascorbate levels on genome-wide gene expression. We selected line L15 (from now on, *vtc2/OE-VTC2*) because it accumulated a higher amount of VTC2-GFP protein than L16. The biomass utilized to determine ascorbate concentration and for RNA extraction comes from the same samples. D) RNA-seq analysis confirms the loss of *VTC2* and *VTC4* expression in their respective T-DNA knockout mutants and shows that the expression of other ascorbate biosynthesis-related genes remains largely unaffected. Different letters in panels C and D denote statistically significant differences (One-Way ANOVA, Tukey post-hoc test, α=0.05). Distributions were tested to meet normality (Shapiro–Wilk’s test) and homoscedasticity (Levene’s test) prior to ANOVA. If not meeting any of these requirements, data was transformed using logarithm and, if meeting the requirements, proceeded to perform ANOVA. Otherwise, Kruskal-Wallis followed by Dunn’s post-hoc test were performed. CBB: Coomassie Brilliant Blue; Glc: glucose, Fru: fructose, Man: mannose, Gal: galactose, GalL: galactono-1,4-lactone, Fuc: fucose; PGI: phosphoglucose isomerase; PMI: phosphomannose isomerase; PMM: phosphomannomutase; GMP: GDP-_D_-mannose pyrophosphorylase (Arabidopsis *VTC1*), GME: GDP-_D_-mannose-3′,5′-isomerase, GGP: GDP-_L_-galactose phosphorylase (*VTC2* and *VTC5*), GPP: _L_-galactose-1-phosphate phosphatase (*VTC4*), _L_-GalDH: _L_-galactose dehydrogenase, _L_-GalLDH: _L_-galactono-1,4-lactone dehydrogenase. FPKM: Fragments Per Kilobase of transcript per Million mapped reads.

Ascorbate concentration is typically higher in leaves and lower in roots, with some species having exceptionally high levels in their fruits (Fenech et al., 2019). In wild-type (WT) Arabidopsis leaves, ascorbate concentration usually ranges from 2 to 5 µmol g^-1^ fresh weight (FW) under low to moderate light intensities, which is comparable to the concentrations of the most prevalent primary metabolites such as sucrose, glucose, serine, and glutamate (Szecowka et al., 2013). However, Arabidopsis *vtc2* mutants with reduced ascorbate levels (∼20% of the WT) remain viable with near to normal growth under benign conditions (Lim et al., 2016). However, ascorbate deficiency results in increased sensitivity to some abiotic stresses and activation of basal pathogen defenses (Barth et al., 2004; Pavet et al., 2005; Mukherjee et al., 2010; Smirnoff and Wheeler, 2024).

To further illuminate the ascorbate’s mechanism of action, we performed a transcriptomic study that leveraged Arabidopsis plants with reduced and elevated ascorbate levels caused by altered expression of *VTC2* and *VTC4* (Fig. 1A). Previous transcriptomic work in Arabidopsis used *vtc2-1* (Kerchev et al., 2011), which carries additional mutations causing small size (Lim et al., 2016) or *vtc1* that also affects the biosynthesis of cell wall precursors (Pastori et al., 2003). Consequently, the transcriptional changes observed reflected both the altered concentration of endogenous ascorbate and structural impairment due to defective growth and cell walls (Lukowitz et al., 2001; Kerchev et al., 2011), making it difficult to disentangle the effects of these two factors. In contrast, the germplasm we chose for our study primarily affects ascorbate biosynthesis. We identified a set of genes whose expression levels correlate with ascorbate concentration and investigated whether a specific transcription factor or subset of transcription factors could regulate gene expression in response to changes in ascorbate levels. In this study, we did not identify specific TFs that correlate with the ascorbate concentration, supporting the notion of complex transcriptional regulation of ascorbate homeostasis in plants, and we characterized the importance of the phytohormone auxin in response to increased ascorbate concentration.

## Results

### The endogenous levels of ascorbate do not exert major control over the expression of genes involved in ascorbate homeostasis

To investigate how different levels of ascorbate affect gene expression, we used Arabidopsis lines with reduced and increased endogenous concentrations of ascorbate relative to the WT. We selected knockout mutant lines of *VTC2*, defective in GGP (*vtc2-4*, from now on *vtc2*; Lim et al., 2016), and of *VTC4,* disrupted in the *L-GALACTOSE 1-PHOSPHATE PHOSPHATASE* (*GPP*) gene (*vtc4-4,* from now on *vtc4*; Torabinejad et al., 2009) (Fig. 1A). We chose these two mutants because *VTC2* and *VTC4* are specifically involved in ascorbate biosynthesis and not in cell wall biosynthesis (Fenech et al., 2019). To obtain lines with elevated ascorbate content, we generated transgenic plants in the *vtc2* background expressing the *VTC2* coding DNA sequence (CDS) lacking the repressive upstream open reading frame (uORF) and fused to *GREEN FLUORESCENCET PROTEIN (GFP)* under the control of the cauliflower mosaic virus (CaMV) *35S* promoter. We have previously shown that *VTC2* is the only gene of the pathway that, upon overexpression, increases ascorbate content (Fenech et al., 2021). Two *vtc2/35S: VTC2_CDS_-GFP* lines were selected, *vtc2/OE-VTC2 L15* and *L16,* that showed higher amounts of VTC2-GFP protein compared with the *vtc2* mutant complemented with the *VTC2p:VTC2_CDS_-GFP* construct driven by the *VTC2* promoter and harboring the repressive uORF (Fenech et al., 2021) (Fig. 1B). For further studies we selected line *L15* (from now on, *vtc2/OE-VTC2*) because it accumulated a higher amount of the VTC2-GFP protein than *L16* (Fig. 1B).

The ascorbate concentration in WT was 3.2 mol g^-1^ FW, while the concentrations for *vtc2, vtc4*, and *vtc2/OE-VTC2* were approximately 20%, 65%, and 165% of WT levels, respectively, indicating a progressive increase in ascorbate concentration among these lines (*vtc2* < *vtc4* < WT < *vtc2/OE-VTC2*) (Fig. 1C). To determine how this range of endogenous ascorbate concentration affects gene expression, we conducted RNA-seq on the same four-week-old WT, *vtc2*, *vtc4*, and *vtc2/OE-VTC2* rosettes from which ascorbate concentration was determined (gene expression and ascorbate concentration are available in Supplementary Material 1). As expected, we found very low expressions of *VTC2* and *VTC4* in their respective mutants (Fig. 1D). Interestingly, altered levels of endogenous ascorbate had very little effect on the expression of genes involved in the mannose/L-galactose ascorbate biosynthesis pathway (Fig. 1D), metabolism, and regulation (Supplementary Figs. S1 and S2). Notably, although the *VTC2* mRNA levels in the CaMV*35S* lines (*vtc2/OE-VTC2*) were lower than those in the line *vtc2/VTC2p:VTC2_CDS_-GFP* where *VTC2* is expressed by its own promoter (Fenech et al., 2021) (Fig. 1D), the protein levels were much higher in the former (Fig. 1B). The elevated VTC2 protein levels in *vtc2/OE-VTC2* lines lacking its regulatory uORF are consistent with the inhibitory role of that uORF on the translation of *VTC2* (Laing et al., 2015).

### Comparative analysis of plants with altered ascorbate content unexpectedly reveals common differentially expressed genes between lines with contrasting levels of ascorbate

We investigated the transcriptional changes in the three Arabidopsis lines with different endogenous ascorbate concentrations (*vtc2, vtc4* and *vtc2/OE-VTC2*) compared with WT. To determine differentially expressed genes (DEGs), we used a false discovery rate (FDR)-corrected *p*-value (*q*-value) <0.05 (Supplementary Material 2). The number of DEGs in the three lines compared to WT (Fig. 2A) and the profile of their volcano plots (Supplementary Fig. S3A) showed a greater difference in the *vtc2* transcriptional program compared to the other lines, supported by the sample outgrouping within the principal component analysis (PCA) plot (Supplementary Fig. S3B). A total of 786 DEGs were found in *vtc2* (506 upregulated and 280 downregulated), 147 DEGs in *vtc4* (123 up and 24 down) and 164 DEGs in *vtc2/OE-VTC2* (135 up and 29 down) (Fig. 2A). These differences in the number of DEGs between *vtc2* and those of *vtc4* and the overexpression line are likely the result of the large difference in ascorbate concentration among these lines (Fig. 1C). We found that ascorbate-deficient mutants *vtc2* and *vtc4* had in common 35 induced genes and 12 repressed genes (Fig. 2B and Supplementary Fig. S4A and B). This non-random overlap (α=0.05, *p*-values of 3.07 x 10^-34^ and 3.12 x 10^-19^ for induced and repressed genes, respectively) was expected since both *vtc2* and *vtc4* have reduced ascorbate levels (Fig. 1C; Supplementary Fig. S4B). Unexpectedly, however, a comparison of DEGs between *vtc4* and *vtc2/OE-VTC2*, the two lines that have reduced or elevated ascorbate content, respectively, shared 82 induced genes (*p*-value=5.28 x 10^-177^) and 4 repressed genes (*p*-value=5.36 x 10^-9^) (Fig. 2B; Supplementary Fig. S4A and B).

**Figure 2.**
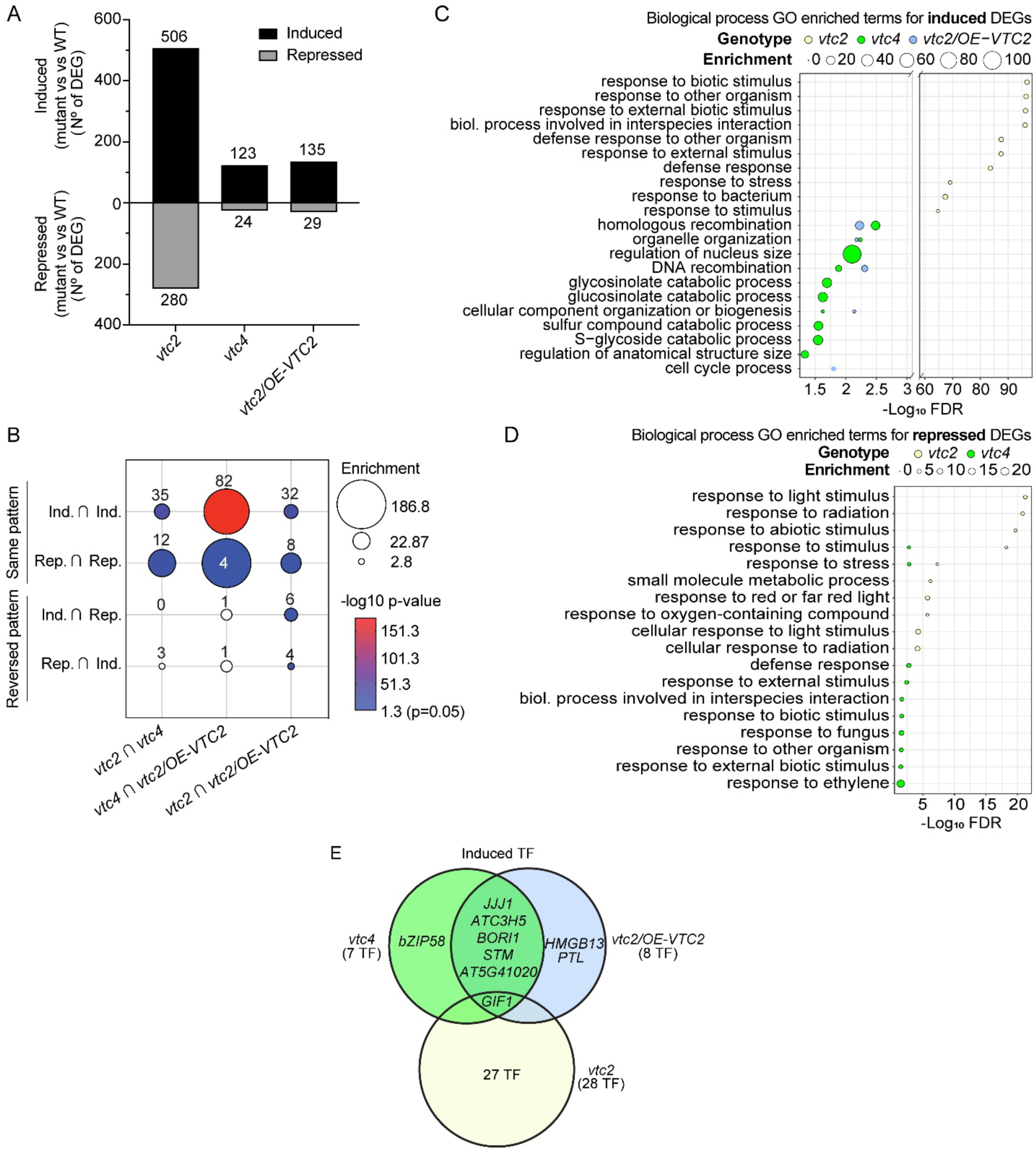
Transcriptome profiles of Arabidopsis ascorbate-altered lines compared to WT. A) Numbers of differentially expressed genes (DEGs) identified at False Discovery Rate (FDR)-corrected p-value (q-value)<0.05. B) Analysis of DEG overlap between the three ascorbate-altered lines. C) GO Biological Process terms associated with induced DEGs. D) GO Biological Process terms associated to repressed DEGs. The symbol ∩ in panel B denotes intersection. E) Overlap of differentially induced TFs between the three ascorbate-altered lines. *bZIP58 (AT1G13600): BASIC LEUCINE-ZIPPER58, JJJ1 (AT1G74250): DNAJ HEAT SHOCK N-TERMINAL DOMAIN-CONTAINING PROTEIN, ATC3H5 (AT1G10320): ZINC FINGER C-X8-C-X5-C-X3-H TYPE FAMILY PROTEIN, BORI1 (AT3G02400): BOREALIN RELATED INTERACTOR 1, STM (AT1G62360): SHOOT MERISTEMLESS, AT5G41020: MYB FAMILY TRANSCRIPTION FACTOR, GIF1 (AT5G28640): GRF1-INTERACTING FACTOR 1*.

To investigate the overlap and differences of their transcriptional programs, we determined the processes impacted by altered ascorbate levels by analyzing the Biological Process Gene Ontology (GO) terms enriched in the induced and repressed DEGs (Fig. 2C and D). The most enriched GO terms for induced DEGs in *vtc2*, but not in *vtc4*, are related to biotic stress, consistent with previous reports where very low ascorbate increased the expression of salicylic acid-responsive genes leading to the higher basal disease resistance of *vtc* mutants (Mukherjee et al., 2010; Kerchev et al., 2011; Pastor et al., 2013) (Fig. 2C). On the other hand, the GO terms enriched for genes repressed in both *vtc2* and *vtc4* partially overlapped for broader terms such as response to stimulus and response to abiotic stress, although *vtc2*-repressed GOs were enriched in abiotic stress and *vtc4*’s in biotic stress (Fig. 2D). Consistent with the overlap between *vtc4-* and *vtc2/OE-VTC2*-induced DEGs, similar enriched GO terms such as homologous recombination, DNA recombination, and organelle organization were identified (Fig. 2C). The reason why both an increase (*vtc2/OE-VTC2*) and a decrease (*vtc4*) of ascorbate concentrations caused a subset of DEGs and GOs terms to respond similarly remains unknown, but it could be the result of an oxidative imbalance of the cell.

### Transcription factors associated with differences between transcriptional programs

We used TDTHub (Grau and Franco-Zorrilla, 2022) and Plant Regulomics (Ran et al., 2020) databases to identify enrichments in DNA motifs and transcription factor (TF) binding sites (TFBSs) within the induced and repressed genes (Supplementary Material 3). The analysis of *vtc2*’s repressed and induced genes revealed a significant enrichment of putative targets from the basic leucine zipper (bZIP) family, like ABA INSENSITIVE 5 (ABI5), and from the WRKY family, respectively, with an FDR<0.05. Among the top ten most enriched TFBS within the promoters of *vtc2*-induced genes, with all of them being the putative targets of WRKY family, we found the recognition sequences of WRKY70, WRKY30, WRKY47, and WRKY55, and these TFs happen to be transcriptionally induced in *vtc2*. The lists of enriched TFBSs in induced DEGs overlapped between *vtc4* and *vtc2/OE-VTC2* but differed from that in *vtc2*. Specifically, *SQUAMOSA PROMOTER BINDING PROTEIN-LIKE3* (*SPL3*/*AT2G33810)* and *MYB DOMAIN PROTEIN116* (*MYB116*/*AT1G25340)* target sites were enriched among both *vtc4-* and *vtc2/OE-VTC2-*induced genes (Supplementary Material 3). For the repressed genes, HOMEOBOX (*AtHB*) TFBSs were enriched among *vtc2/OE-VTC2-*downregulated genes, but no enrichment was found within *vtc4-* downregulated genes by either TDTHub or Plant Regulomics.

To identify DEGs encoding transcriptional regulators (TRs), i.e., TFs and other regulators not binding DNA, that may account for the differences in the transcriptomic profiles of the different ascorbate lines, we used the list of manually curated Arabidopsis TRs obtained from AGRIS (Davuluri et al., 2003; Palaniswamy et al., 2006; Yilmaz et al., 2011; Spurney et al., 2020) (Supplementary Material 4). A total of 50 TR genes (28 upregulated, 22 downregulated) were identified among *vtc2*-dependent DEGs, while 7 and 8 TRs were identified among *vtc4-* and *vtc2/OE-VTC2-*controlled DEGs, respectively, with all of them induced. Remarkably, the *vtc4-* and *vtc2/OE-VTC2-*regulated TR gene lists shared 6 upregulated genes, including *SHOOT MERISTEMLESS* (*STM*) involved in the maintenance of shoot and flower meristem (Endrizzi et al., 1996) (Fig. 2E). Interestingly, all lines shared one differentially expressed TR gene, *GROWTH REGULATING FACTOR1 (GRF1)-INTERACTING FACTOR1/ANGUSTIFOLIA3* (*GIF1/AN3/AT5G28640*), which coordinates leaf meristem positioning, cell proliferation, and postmitotic cell expansion in leaves (Kawade et al., 2010) (Fig. 2E). Notably, among the TRs upregulated in *vtc2*, 18 out of 28 genes belong to the *WRKY* (10 genes) and *NO APICAL MERISTEM/ARABIDOPSIS TRANSCRIPTION ACTIVATION FACTOR/CUP-SHAPED COTYLEDON* (*NAC*) (8 genes) families (Fig. 2E), with many of them implicated in the response to biotic stress, crosstalk with ROS, hormone signaling, and leaf senescence (Chen et al., 2019; Yuan et al., 2019; Bian et al., 2020; Ahmad et al., 2024). In contrast, some *ETHYLENE RESPONSE FACTOR* (*ERF*) genes like *ERF018/ACTIVATOR PROTEIN BINDING TO CMSRE-1 3 (ACRE3), ERF094/ OCTADECANOID-RESPONSIVE AP2/ERF DOMAIN59 (ORA59*), and *ERF023* were among the most-repressed TR genes in *vtc2* (Supplementary Material 4). In addition, TRs associated with shade avoidance response and response to nitrogen concentration by modulating cytokinin (*BRASSINOSTEROID-ENHANCED EXPRESSION1* (*BEE1)* (Gautrat et al., 2024) and *CYTOKININ RESPONSE FACTOR2* (*CRF2)* (Chen et al., 2024)) and auxin (*IAA6/SHORT HYPOCOTYL1* (*IAA6/SHY1)* (Kim et al., 1996; Nguyen et al., 2023) and *IAA17/AUXIN RESISTANT 3* (*IAA17/AXR3*) (Leyser et al., 1996)) signaling pathways were also repressed in *vtc2*. These results suggest an interplay between ascorbate and hormone signaling.

### Refining the regulatory landscape of ascorbate-correlated genes

Variations in ascorbate concentration across the analyzed samples enabled the identification of genes whose expression correlated with endogenous ascorbate levels. Using Pearson correlation analysis with an FDR-corrected p-value threshold (q-value<0.05), we identified 474 positively correlated genes (PCGs) and 376 negatively correlated genes (NCGs) (Supplementary Material 5). Further filtering based on differential expression between *vtc2* and *vtc2/OE-VTC2* (t-test, q-value<0.05) narrowed this down to 90 PCGs and 107 NCGs whose expression in *vtc2* is complemented by *vtc2/OE-VTC2* (Fig. 3A and B; Supplementary Material 6A and B). The expression patterns of these genes visually confirmed their correlation with ascorbate concentration (Fig. 3C and D).

**Figure 3.**
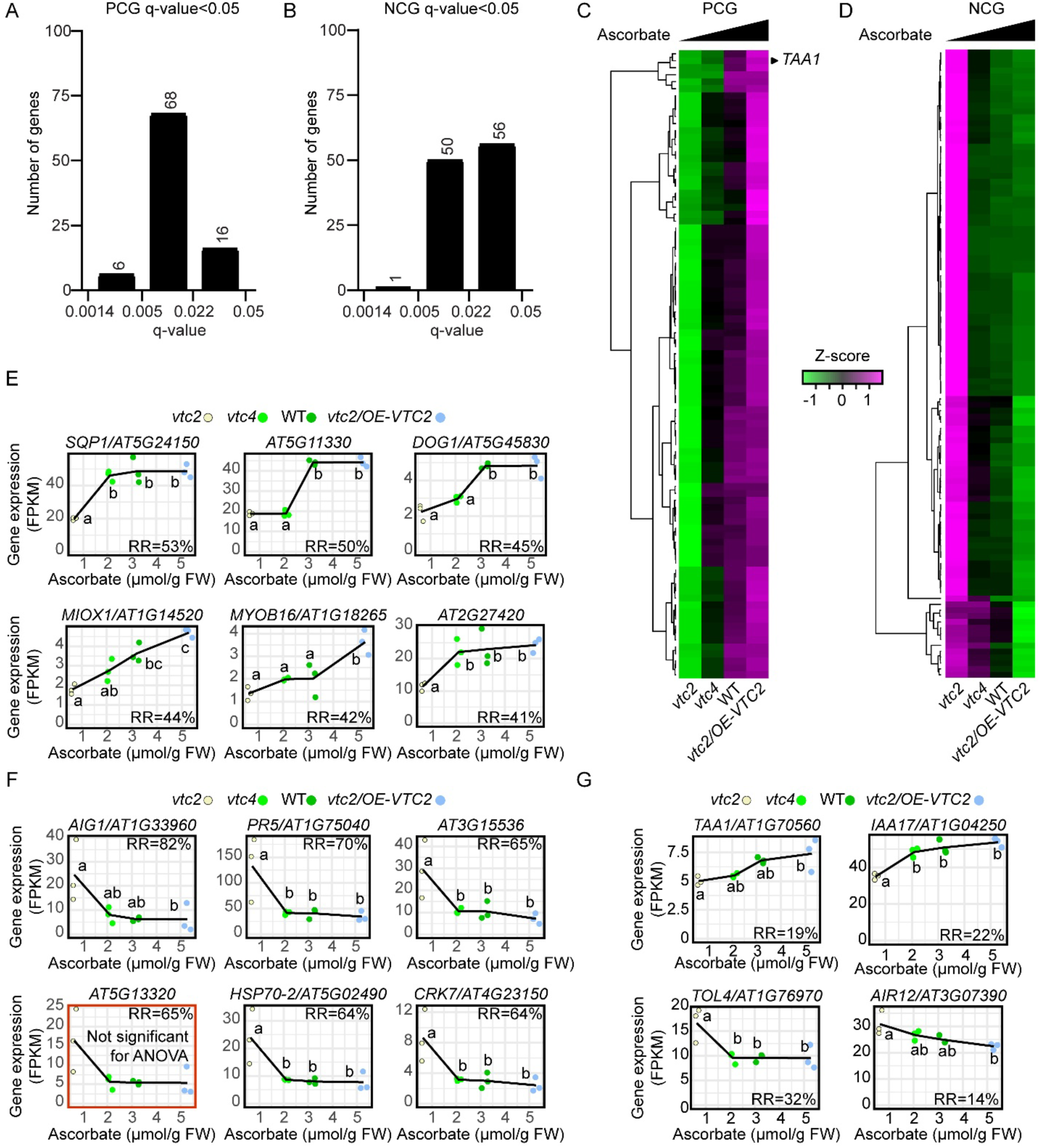
Identification of genes whose expression is correlated with ascorbate concentrations across genotypes and that are differentially expressed between the most contrasting ascorbate-altered lines, *vtc2* and *vtc2/OE-VTC2*. A,B) Distribution of statistical confidence values for genes positively (A) and negatively (B) correlated with ascorbate concentrations for genes with a False Discovery Rate (FDR)-corrected p-value (q-value)<0.05 cutoff. C, D) Expression profiles of positively correlated genes (PCGs) (C) and negatively correlated genes (NCGs). (D) Heatmap generated with iDEP (Ge et al., 2018). Hierarchical clustering of PCGs and NCGs was calculated by Pearson correlation distances. The Z-score normalization of the expression data (FPKM) was performed by subtracting the average expression value for each gene and dividing by the standard deviation. E) The six strongest PCGs based on their relative response (RR) calculated as explained in Supplementary Fig. S6. F) The six strongest NCGs based on their relative response (RR) calculated as explained in Supplementary Fig. S6. G) Genes with prominent roles in auxin homeostasis that showed significant correlation with ascorbate concentration. RNA-seq FPKM values (y-axes) vs ascorbate concentrations (x-axes) are plotted. Different letters denote statistically significant differences (One-Way ANOVA, Tukey post-hoc test, α=0.05). Distributions were tested to meet normality (Shapiro–Wilk’s test) and homoscedasticity (Levene’s test) prior to ANOVA. If not meeting any of these requirements, data was transformed using logarithm and, if meeting the requirements, proceeded to perform ANOVA. Otherwise, Kruskal-Wallis followed by Dunn’s post-hoc test were performed. FPKM: Fragments Per Kilobase of transcript per Million mapped reads, FW: fresh weight; *SQP1: SQUALENE MONOOXYGENASE, AT5G11330: FAD/NAD(P)-BINDING OXIDOREDUCTASE*, *DOG1: DELAY OF GERMINATION 1, MIOX1; MYO-INOSITOL OXYGENASE1, MYOB16: MYOSIN BINDING PROTEIN 16, AT2G27420: CYSTEINE PROTEINASES, AIG1: AVRRPT2-INDUCED GENE 1, PR5: PATHOGENESIS-RELATED GENE 5, AT3G15536: Unknown gene, AT5G13320: AVRPPHB SUSCEPTIBLE 3, HSP70-2: HEAT SHOCK 70 PROTEIN, CRK7: CYSTEINE-RICH RECEPTOR-LIKE PROTEIN KINASE 7, TAA1: TRYPTOPHAN AMINOTRANSFERASE OF ARABIDOPSIS 1, IAA17/AXR3: AUXIN RESISTANT 3/INDOLE-3-ACETIC ACID INDUCIBLE 17, TOL4: TOM1-LIKE 4, AIR12: AUXIN-INDUCED IN ROOT CULTURES 12*.

To better characterize their response to varying ascorbate levels, we examined expression fold changes in samples with the most contrasting ascorbate concentrations—*vtc2* (low ascorbate) and *vtc2/OE-VTC2* (high ascorbate) (Supplementary Fig. S5A and B). Genes were classified into three categories: (i) those responsive to ascorbate depletion (44 PCGs, 57 NCGs), (ii) those responsive to ascorbate excess (25 PCGs, 35 NCGs), and (iii) those exhibiting a proportional response to ascorbate levels (expected response; 21 PCGs, 15 NCGs). To quantify their relative response, we assessed the fold-change distance from the origin (0,0) in Supplementary Fig. S5A,B, normalized against a theoretical 1:1 response between fold change in gene expression and ascorbate levels (Supplementary Fig. S6A and B). PCGs primarily showed mild responses (12.5-37.5% of an ideal PCG), with only one gene surpassing 50%. In contrast, a greater proportion of NCGs exceeded 50%, with some reaching ∼80% (Supplementary Fig. S6C). These findings suggest that transcriptional changes due to ascorbate fluctuations may be mediated by additional factors such as reactive oxygen species (ROS) or other signaling molecules.

### Transcriptional Regulation of Ascorbate-Correlated Genes

To investigate whether ascorbate-correlated genes are co-regulated by specific transcription factors (TFs), we leveraged the DAP-seq dataset (O’Malley et al., 2016). We focused on TFs targeting the promoters of PCGs and NCGs responding proportionally to ascorbate levels. Hierarchical clustering of PCG and NCG based on the TF that they are targeted by, and spatially arranged based on their relative response (using their FC *vtc2*), allowed us to identify distinct TF subsets (Supplementary Fig. S5C and D; Supplementary Material 7). Among PCGs, circadian clock-associated TFs, including REVEILLE4 (RVE4), RVE8, and LATE ELONGATED HYPOCOTYL (LHY), as well as ERFs were prominent (Supplementary Fig. S5C; Supplementary Material 7A). In contrast, NCGs were primarily targeted by WRKY TFs such as WRKY18, WRKY33, and WRKY40, known for their roles in biotic stress responses (Liu et al., 2015; Birkenbihl et al., 2017) (Supplementary Fig. S5D; Supplementary Material 7B). Remarkably, these TFs were not specifically targeting the “responding as expected” class of NCGs, and many other correlated genes were also targeted by WRKYs (Supplementary Material 8), suggesting that they may play a broader role in ascorbate-mediated transcriptional regulation. To further explore the possibility that a subset of TF could regulate the expression of ascorbate-correlated genes, we arranged the TF-target interaction based on the relative response of correlated genes, i.e., the fold-change difference in expression compared to the fold-change difference in ascorbate concentration (Supplementary Fig. S6), regardless of their type of response (depletion, expected, or excess). Unfortunately, we could not identify any TF subset that could explain a stronger response to ascorbate concentration (Supplementary Material 9), suggesting that the observed changes in gene expression cannot be explained solely by the combination of TF targeting these genes, and that additional layers of regulation may be operating in the ascorbate-mediated control of gene expression.

### Visualizing Gene Sensitivity to Ascorbate Depletion

Given the intermediate ascorbate deficiency in *vtc4* mutants, we explored gene expression patterns across *vtc4* and *vtc2* to assess gene expression sensitivity to ascorbate depletion. By comparing fold changes in *vtc4* (65% of WT ascorbate) *versus vtc2* (20% of WT), we classified genes based on their depletion sensitivity (Supplementary Fig. S7A and B). Statistical analysis revealed significant enrichment of genes with low or expected sensitivity to ascorbate deficiency (observed 66%, expected 20%), while high-sensitivity genes were underrepresented (observed 33%, expected 79.9%) (Supplementary Fig. S7C and D). Similarly, PCGs and NCGs responding to ascorbate depletion were enriched more than threefold (Supplementary Fig. S7C and D). Considering that *vtc2-4* mutant does not show obvious developmental defects despite containing 20% of WT content (Lim et al., 2016), and that some Arabidopsis ecotypes have ascorbate concentrations that fall between the levels of Col-0 *vtc2* and Col-0 *vtc4* even under normal growth conditions (Aarabi et al., 2023), the selective conservation of low-sensitivity genes may suggest an evolutionary preference for energy-efficient transcriptional responses unless critical for survival.

Interestingly, as observed in Supplementary Figs. S5A and B and S7A and B, genes with similar ascorbate sensitivity did not necessarily share the same type of response. To integrate these variables, we developed a custom plot (Supplementary Fig. S8A and B), simultaneously visualizing gene response type (y-axis) and sensitivity to ascorbate depletion (x-axis). Expected responses aligned near the red horizontal line, and expected sensitivities aligned near the blue vertical line. Therefore, genes highly sensitive to depletion positioned towards the right of the vertical blue line (Supplementary Fig. S8C, sectors ii and iv). For example, PCGs like *MYO-INOSITOL OXYGENASE1* (*MIOX1*), *TAA1*, and *PLANT U-BOX48* (*PUB48*) exhibited high sensitivity, while most NCGs clustered in sector (i), characterized by an upregulation in *vtc2* but negligible changes elsewhere (Supplementary Fig. S8B and D). Notably, nine PCGs exhibited expected responses and high sensitivity (Supplementary Fig. S9A; see green highlighted genes in Supplementary Material 6A). Conversely, only three NCGs met similar criteria (Supplementary Fig. S9B; see green highlighted genes in Supplementary Material 6B). These genes had not previously been linked to ascorbate, most likely due to the weak change of expression between the two most contrasting genotypes, *vtc2* and *vtc2/OE-VTC2,* with relative responses ranging between 11-19% (Supplementary Fig. S9). However, genes with stronger relative responses like *DELAY OF GERMINATION1 (DOG1)* (PCG; Fig. 3E; see blue highlighted genes in Supplementary Material 6A), or *AvrRpt2-INDUCED GENE1 (AIG1), CYSTEINE-RICH RECEPTOR-LIKE PROTEIN KINASE7 (CRK7)* and *PATHOGENESIS-RELATED GENE5 (PR5)* (NCG; Fig. 3F; see blue highlighted genes in Supplementary Material 6B) are some examples of genes whose change in expression has been recapitulated in other studies of the *vtc2-1* mutant (Gao et al., 2011; Kerchev et al., 2011).

In summary, Supplementary Fig. S8 provides a refined framework for identifying genes with distinct transcriptional responses to ascorbate (Supplementary Material 6A and B). Our findings provide a foundation for future research into ascorbate-mediated gene regulation and its implications for redox homeostasis in plant cells.

### Expression patterns of PCGs and NCGs reveal a potential link between ascorbate and auxin

Among correlated genes, we identified several PCGs and NCGs involved in different aspects of auxin homeostasis, which is in line with prior studies that suggested that there might be an interaction between ascorbate and auxin in plants (Akram et al., 2017; Bulley et al., 2021; Wang et al., 2021). Among the PCGs of the class “responding to ascorbate as expected” that also showed “high sensitivity to depletion”, we found *TAA1*, a tryptophan aminotransferase catalyzing the first committed step of auxin biosynthesis (Stepanova et al., 2008; Tao et al., 2008; Yamada et al., 2009), whose expression is reduced in *vtc2* and *vtc4*, and increased in the *vtc2/OE-VTC2* line (Fig. 3C and G; Supplementary Figs. S8A and S9A). While other genes involved in auxin biosynthesis were either very poorly expressed or not significantly correlated with ascorbate concentration, a *TAA1* paralog, *TAR2,* was among genes repressed in *vtc2* but not significantly correlated with ascorbate concentration (Supplementary Fig. S10; Supplementary Material 2). In addition, a number of genes involved in auxin signaling and response were also identified among the ascorbate co-regulated genes (Fig. 3G). For example, *IAA17/AXR3,* that encodes an auxin co-receptor (Leyser et al., 1996; Kubalová et al., 2024), and *ROOT MERISTEM GROWTH FACTOR9* (*RGF9),* encoding a peptide involved in the subcellular regulation of auxin carrier turnover (Whitford et al., 2012), are both PCGs responding to ascorbate depletion (Fig. 3G; Supplementary Figs. S8A and S9A; Supplementary Material 6A). Among NCGs, we identified those encoding the ubiquitin-binding protein *TOBAMOVIRUS MULTIPLICATION1 (TOM1)-LIKE4* (*TOL4*) involved in subcellular distribution of PIN-type auxin carriers (Korbei et al., 2013) (Fig. 3G) and *AUXIN-INDUCED IN ROOT CULTURES12* (*AIR12*), an auxin-inducible gene (Neuteboom et al., 1999) that provides cold tolerance by increasing ascorbate concentration, hence counteracting H_2_O_2_ accumulation (Wang et al., 2021) (Fig. 3G; Supplementary Fig. S8B; Supplementary Material 6B). In summary, the changes in expression of genes related to auxin biosynthesis, transport, and signaling point towards a potential crosstalk between ascorbate concentration and auxin patterns of activity.

### Exogenous ascorbate induces auxin biosynthesis under continuous light, but not under continuous darkness

Encouraged by prior studies suggesting that ascorbate and auxin may interact (Kerk et al., 2000; Bulley et al., 2021; Wang et al., 2021), and our finding that several genes involved in auxin biosynthesis, signaling, or transport are correlated with ascorbate concentration, we decided to further investigate the relationship between the auxin and ascorbate levels. We focused on *TAA1*/AT1G70560 (also known as *WEAK ETHYLENE INSENSITIVE8* (*WEI8*)) because it is a key component in auxin biosynthesis implicated in the crosstalk with several hormones and other signals (Stepanova et al., 2008; Tao et al., 2008; Yamada et al., 2009; Zhou et al., 2011; Brumos et al., 2018; Brumos et al., 2020). We chose to perform this work in seedlings (rather than in adult plants which we used for RNA-seq) given that most studies cited above exploring plant defects in auxin biosynthesis or response also looked at seedlings, and we relied on these established phenotypic assays as our reference. We analyzed whether altered ascorbate concentration affects auxin biosynthesis through *TAA1* expression by examining the effect of exogenous application of ascorbate on the expression of *TAA1* using the recombineering line *wei8-2*/*TAA1p:YPet-gTAA1* (backcrossed to WT, previously shown in *wei8-2 tar2-1* background in Brumos et al., 2018). In this line, the *TAA1* promoter (10 kb upstream of the ATG start codon) drives the expression of the yellow fluorescence protein gene *YPet* fused to the N-terminus of the full-length *TAA1* gene (containing all introns) in its genomic context, inserted in a random location of the *wei8-2* mutant genome (*wei8*/*YPet-gTAA1*). Consistent with the transcriptomic data, the exogenous application of ascorbate to *wei8*/*YPet-gTAA1* seedlings caused the induction of the YPet-TAA1 fusion protein in the epidermis and vasculature in the transition and elongation zones of the root in plants grown under continuous light (Fig. 4A). To determine whether the application of ascorbate also led to increased auxin signaling, we used Arabidopsis lines expressing the synthetic auxin-inducible reporter *DR5:GFP* in WT (Col-0) background (Ulmasov et al., 1997; Weijers et al., 2006). Under control conditions, GFP was detected in the root columella, quiescent center, and vasculature cells. Upon ascorbate supplementation, *DR5:GFP* expression increased in the non-meristematic portion of the root vasculature (Fig. 4B).

**Figure 4.**
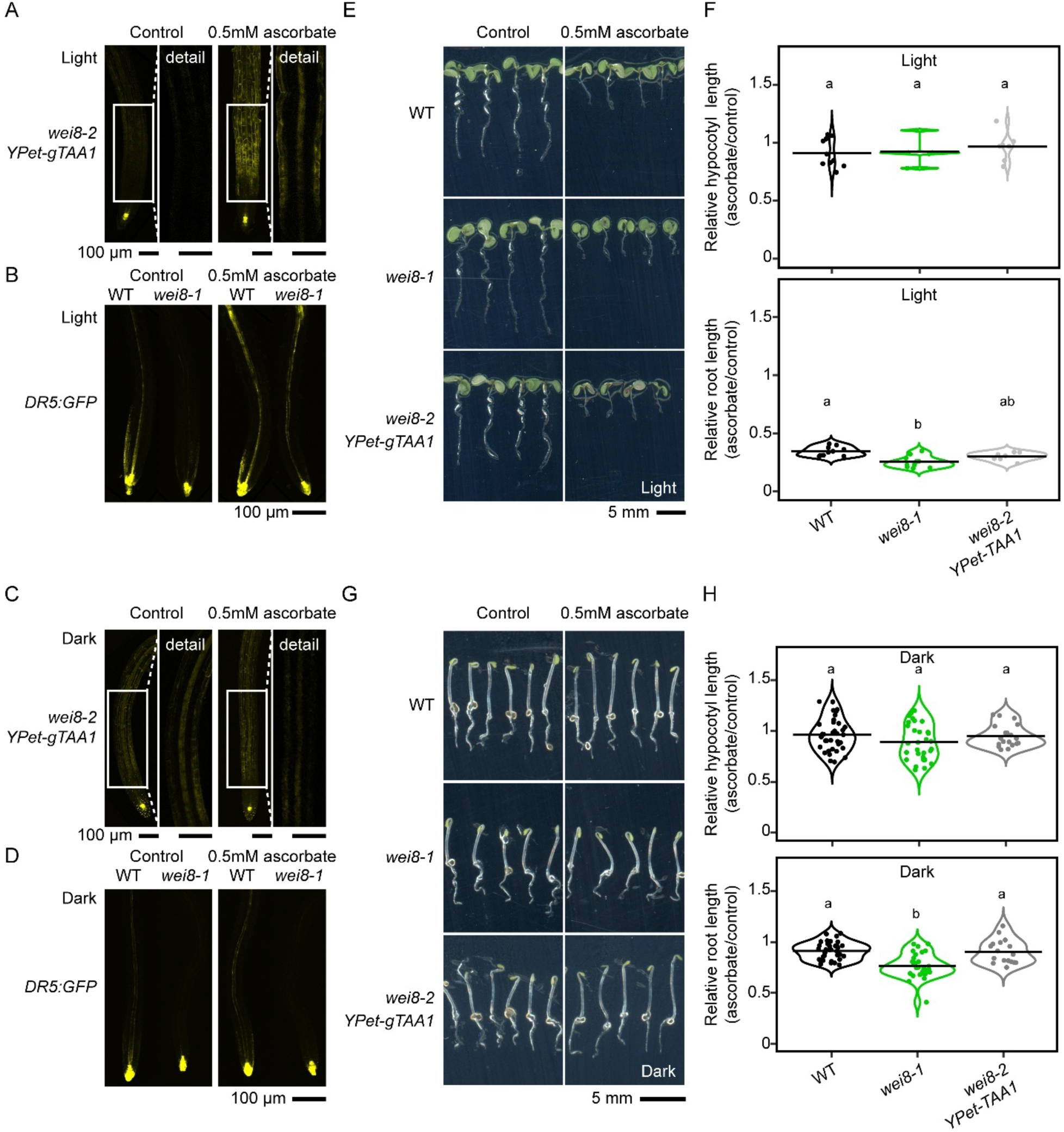
Ascorbate induces auxin biosynthesis in Arabidopsis in a light-dependent manner. Panels A, B, E, and F show seedlings grown for 5 days under continuous LED light and panels C, D, G, and H show seedlings grown for 3 days in the dark. A, C) Expression of a recombineering genomic DNA construct, *TAA1p:YPet-gTAA1*, in *wei8-2* roots is increased in light-but not dark-grown seedlings germinated in 0.5 mM ascorbate. B, D) Expression of an auxin-inducible reporter, *DR5:GFP,* in WT and *wei8* mutant backgrounds is induced in roots in light-but not dark-grown seedlings germinated in 0.5 mM ascorbate. E, F) The length of roots, but not that of hypocotyls, of light-grown WT seedlings germinated in the media supplemented with 0.5 mM ascorbate is reduced to 35% of the untreated seedlings’ length, with *wei8* showing ascorbate hypersensitivity in roots (25% of untreated roots’ length). G, H) The length of WT roots and hypocotyls of dark-grown seedlings exposed to 0.5 mM ascorbate is statistically indistinguishable from that of untreated plants (90%), whereas *wei8* displays a 25% root growth inhibition in ascorbate. Different letters denote statistically significant differences (One-Way ANOVA, Tukey post-hoc test, α=0.05). Distributions were tested to meet normality (Shapiro–Wilk’s test) and homoscedasticity (Levene’s test) prior to ANOVA. If not meeting any of these requirements, data was transformed using logarithm and, if meeting the requirements, proceeded to perform ANOVA. Otherwise, Kruskal-Wallis followed by Dunn’s post-hoc test were performed. More than 20 roots per genotype and treatment have been scored through experimental repetitions and a representative image was selected. All microscopy images are Z stacks through the root from the cortical to the equatorial plane, except the details shown in panels A and C, that are stacks of 2-5 equatorial planes.

We examined whether the *DR5:GFP* expression changes in response to ascorbate in the absence of functional *TAA1*. The *wei8-1* mutant (hereon, *wei8*), whose tryptophan aminotransferase activity encoded by *TAA1* has been knocked-out, showed an overall reduction of *DR5:GFP* signal in control conditions relative to WT, consistent with partially impaired auxin biosynthesis in the mutant (Fig. 4B). However, ascorbate treatment still caused an increase in *DR5* activity in the vasculature in the *wei8 DR5:GFP* line (Fig. 4B) suggesting that ascorbate may regulate the levels and distribution of auxin even in the absence of functional *TAA1*. In contrast to what was seen in seedlings germinated under continuous light, in dark-grown seedlings, the YPet-TAA1 expression and the *DR5:GFP* activity in roots was not affected by exogenous ascorbate (Fig. 4C and D), suggesting that the effect of ascorbate on auxin response in roots may be light-dependent. In light-grown WT seedlings treated with 0.5mM ascorbate, a ∼65% root growth reduction was observed relative to the untreated control, whereas the ascorbate response of hypocotyls was negligible (Fig. 4E and F). Consistent with unaltered levels of auxin activity in the dark, and in contrast to light-grown seedlings, the growth of seedlings in the dark was not prominently affected by the supplementation of growth media with 0.5mM ascorbate (Fig. 4G and H). Importantly, regardless of light conditions, the reduction of root growth by ascorbate was significantly more prominent in the *wei8* mutant, and the response of roots was reverted to WT values in the complemented line, *wei8*/*YPet-gTAA1* (Fig. 4C, D, G, H). The greater sensitivity to exogenous ascorbate observed in the primary roots of *wei8* mutants was consistently reproduced at lower concentrations of ascorbate in light-grown seedlings (Supplementary Fig. S11). On the other hand, hypocotyl response to ascorbate was unaffected by the genotype in both sets of conditions (Fig. 4C, D, G, H), with enhanced hypocotyl elongation known to be associated with increased auxin activity under standard growth conditions (*superroot2* mutant in Delarue et al., 1998; *YUC1* overexpression in Zhao et al., 2001; *TAA1p:YUC1* transgene in Stepanova et al., 2011; *iaaM* in Romano et al., 1995; and *cyp72ox* in Zhao et al., 2002). In summary, we conclude that exogenously applied ascorbate in light-grown seedlings translated into an increase in TAA1 protein accumulation and enhanced auxin response in specific cell types of the primary root.

### *TAA1* and *TAR2* have a prominent role in the light-dependent ascorbate-mediated increase in auxin response

To investigate ascorbate-dependent induction of a wider set of auxin biosynthesis genes beyond *TAA1*, we analyzed the expression of its two paralogs, *TAR1* and *TAR2*, and 11 *YUC* genes in response to ascorbate treatment by utilizing Arabidopsis recombineering lines that harbor genomic DNA fusions with *GUS,* and in parallel we employed a *DR5:GUS* reporter to monitor auxin activity (Stepanova et al., 2008; Brumos et al., 2020). Consistent with the results presented above for the *DR5:GFP* reporter (Fig. 4B), in light-grown seedlings, ascorbate supplementation of growth media led to the expansion of the domain of *DR5:GUS* expression in the root vasculature, but not in the hypocotyls (Fig. 5A and B). Likewise, the pattern of *GUS-gTAA1* activity in roots upon ascorbate application in light-grown plants (Fig. 5A) was similar to that observed in the *YPet-gTAA1* line (Fig. 4A), with increased GUS staining in the root elongation zone. Interestingly, in light-grown seedlings, ascorbate treatment also induced the expression of *GUS-gTAR2* (Fig. 5A), a gene encoding the same enzymatic activity as *TAA1* (Stepanova et al., 2008), which could explain the ascorbate-dependent increase of auxin activity in the *wei8* mutant (Fig. 4B). Remarkably, the induction of *TAR2* expression, like that of the *TAA1* and *DR5* reporter lines, was also light-dependent, i.e. not observed in dark-grown seedlings (Fig. 5B). We also found that ascorbate application altered the expression of other auxin biosynthetic genes, such as *YUC5*, *YUC6*, and *YUC8,* in light-grown seedlings (Supplementary Fig. S12A), while only *YUC3* was affected by ascorbate in the dark (Supplementary Fig. S12B). While these results may appear not fully supported by our transcriptomic analysis that failed to detect the effects of endogenous ascorbate levels on *YUC* gene expression (Supplementary Fig. S7), they are not unexpected given the differences in plant age, growth conditions, and tissue types between these experiments and our RNA-seq work. Furthermore, the ascorbate-triggered expression domain changes visualized with the help of auxin reporters are highly spatially restricted and thus are likely to be diluted in samples comprising whole plants analyzed by RNA-seq. Finally, the reporter lines examined in this work harbor translational fusions that could be subject to translational and post-translational regulations not accounted for at the mRNA level.

**Figure 5.**
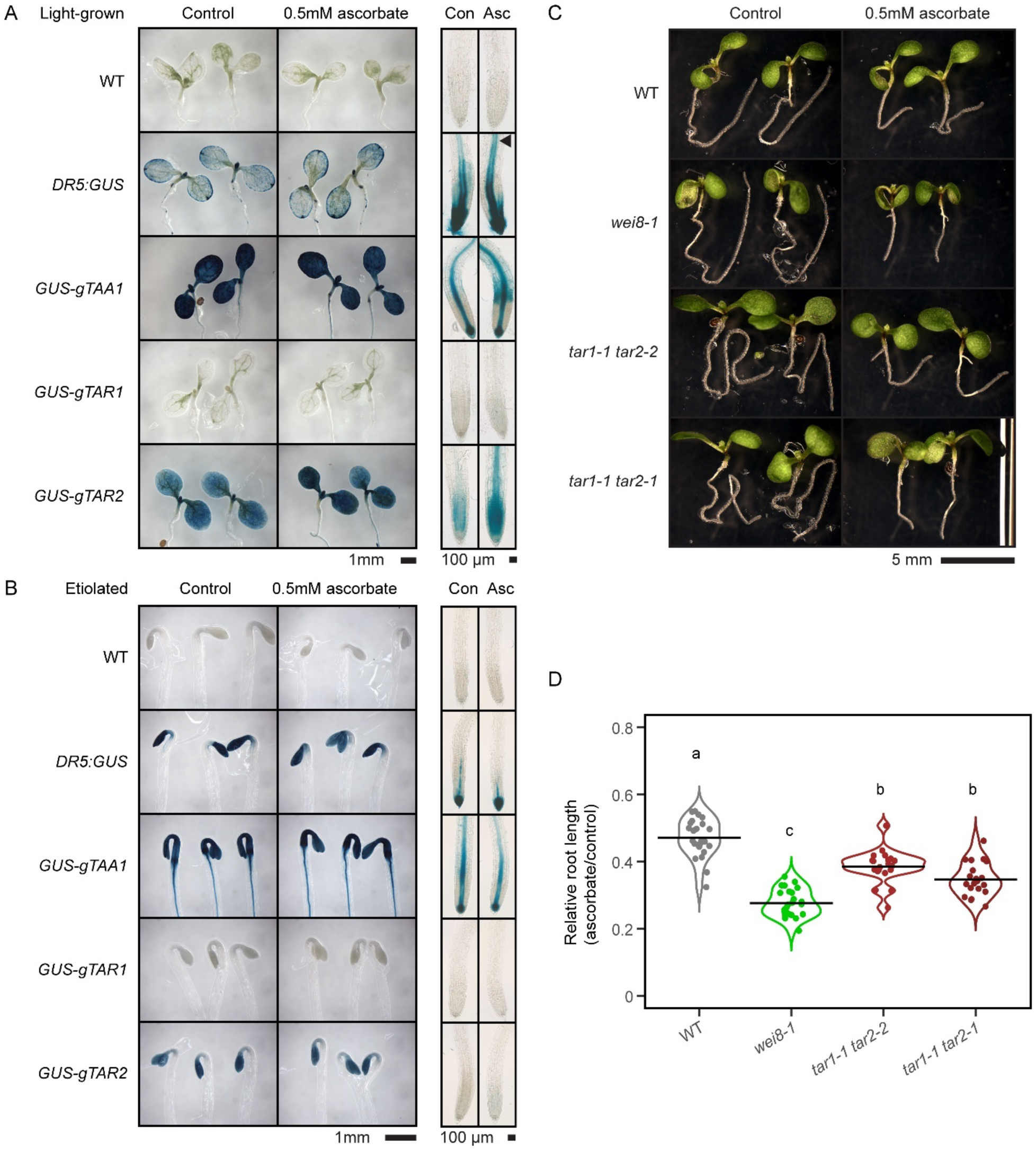
Ascorbate (Asc)-induced auxin biosynthesis in Arabidopsis relies on *TAA1* and *TAR2*. Panels A, C and D show seedlings grown for 5 days under continuous light, and panel B shows seedlings grown for 3 days in the dark. A, B) Auxin-inducible reporter *DR5:GUS* and recombineering-generated translational fusions of *TAA1* and *TAR2* with *GUS* show increased reporter activity upon media supplementation with 0.5mM ascorbate in light-but not dark-grown plants. C, D) Plants exposed to 0.5mM ascorbate in the media display reduced root growth as compared to root elongation in the absence of ascorbate. *wei8* mutant shows greater sensitivity to 0.5 mM ascorbate than WT, *tar1 tar2-2* and *tar1 tar2-1,* with *tar1 tar2-2* and *tar1 tar2-1* showing intermediate response between that of *wei8* and WT. Different letters denote statistically significant differences (One-Way ANOVA, Tukey post-hoc test, α=0.05). Distributions were tested to meet normality (Shapiro–Wilk’s test) and homoscedasticity (Levene’s test) prior to ANOVA. If not meeting any of these requirements, data was transformed using logarithm and, if meeting the requirements, proceeded to perform ANOVA. Otherwise, Kruskal-Wallis followed by Dunn’s post-hoc test was performed.

Considering the prominent ascorbate-induced expression of *TAA1* and *TAR2*, the two major active tryptophan aminotransferase genes, we tested the root growth response to ascorbate of a *wei8-1 tar2-1* mutant. However, this double mutant shows a severe reduction in root growth in control conditions, consistent with a severely compromised tryptophan aminotransferase activity (Stepanova et al., 2008) and appears to have a more severe response to exogenous ascorbate (Supplementary Fig. S13). We could not reliably quantify this response given the need to work with a segregating *wei8 tar2/+* population as *wei8 tar2* double homozygotes are fully sterile (Stepanova et al., 2008). To further confirm that *TAA1* and *TAR2* genes are the only genes with prominent tryptophan aminotransferase activity implicated in ascorbate-induced auxin biosynthesis, we characterized the response of the WT, *wei8,* and two different *tar1 tar2* mutants (a stronger *tar1 tar2-*1, and a weaker *tar1 tar2-2*) upon ascorbate supplementation of light-exposed seedlings (Stepanova et al., 2008). While all four genotypes showed similar organ growth in control conditions, *wei8* and both *tar1 tar2* double mutants were hypersensitive to ascorbate in the roots (Fig. 5C and D). Interestingly, the two *tar1 tar2* mutant allele combinations showed an intermediate FC in root length in response to ascorbate that fell between that of WT and *wei8* (Fig. 5C and D). A mild disruption of *TAR2* (*tar1 tar2-2*) lead to phenotypes that are slightly more similar to WT, whereas a stronger disruption of *TAR2* (*tar1 tar2-1*) translated into a root length FC more similar to that of *wei8* (Fig. 5C and D). This result, together with our observation that *TAR2*, but not *TAR1*, is expressed in the roots under continuous light and it is induced upon ascorbate exogenous supplementation (Fig. 5A), suggests that both *TAA1* and *TAR2* play a role in the root response to ascorbate. The fact that *wei8* and *tar1 tar2* mutants showed a more severe reduction of root growth after ascorbate treatment than WT implies that auxin is necessary to counteract ascorbate-induced root shortening. Consistently, ascorbate induced the expression of *TAA1* and *TAR2* (Figs. 4A and 5A), leading to an increase in auxin activity.

### Tissue-specific auxin gradients are necessary to cope with an increase in ascorbate concentration

In light of these results, we tested whether exogenous supplementation with IAA could improve root growth of WT and auxin-deficient seedlings in the presence of ascorbate. To do that, we analyzed the relative root length of WT, *wei8,* and *aux1-7* (hereon, *aux1*, an auxin-insensitive mutant defective in IAA uptake into the cells; Bennett et al., 1996) in light-grown seedlings germinated in the presence of 0.5 mM ascorbate either with or without supplementation with 100 nM IAA (Fig. 6A and B). We observed that the root lengths of WT, *wei8* and *aux1* plants in control conditions (no ascorbate) remained unchanged in the presence of 100 nM IAA (Fig. 6A). However, we found that 100 nM IAA partially alleviated the inhibitory effect of 0.5mM ascorbate on WT root growth (Fig. 6B). This effect was also observed in the mild auxin-deficient mutant *wei8* but not in the strong auxin mutant, *aux1*. Interestingly, the relative root length in IAA plus ascorbate in *wei8* was shorter than that of WT, suggesting that the low auxin levels of *wei8* may not be the only reason for the short root phenotype of this mutant in the presence of ascorbate. We reason that exogenously applied IAA may not result in the same IAA distribution patterns or gradients as those produced by the combination of *TAA1*-mediated local auxin production in response to ascorbate and auxin transport.

**Figure 6.**
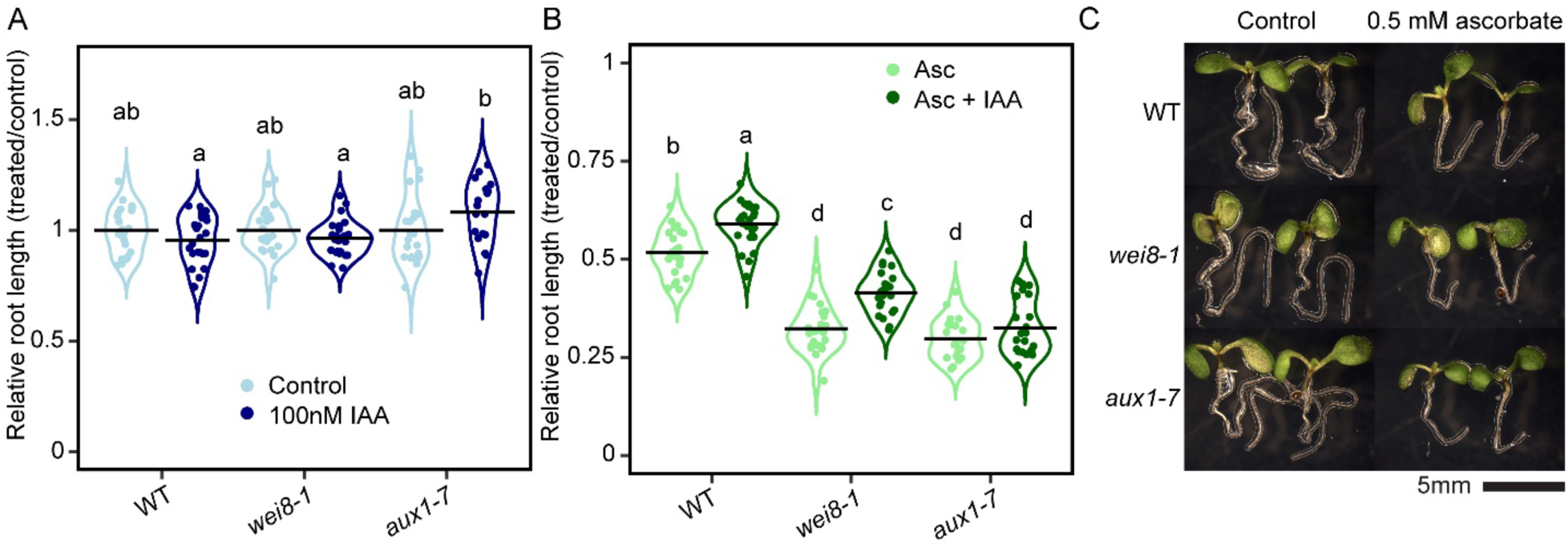
Exogenous auxin (IAA) partially reverts the root hypersensitivity to ascorbate in *wei8*, but not in *aux1*. Seedlings were germinated for 5 days under continuous light. A) Presence of 100 nM IAA in growth media does not inhibit root growth. B) 100 nM IAA enhances the root growth in presence of 0.5 mM ascorbate (Asc), but it does not fully restore *wei8* and *aux1* mutants’ primary root growth. C) Representative images of the phenotypes quantified in A and B. Different letters denote statistically significant differences (One-Way ANOVA, Tukey post-hoc test, α=0.05). Distributions were tested to meet normality (Shapiro–Wilk’s test) and homoscedasticity (Levene’s test) prior to ANOVA. If not meeting any of these requirements, data was transformed using logarithm and, if meeting the requirements, proceeded to perform ANOVA. Otherwise, Kruskal-Wallis followed by Dunn’s post-hoc test was performed. IAA: indole-3-acetic acid

It is known that IAA enters the cells mainly by auxin influx carriers like AUX1 and LAXs (Swarup and Bhosale, 2019). To investigate the potential role of auxin influx in the root response to ascorbate, we analyzed the relative root growth inhibition caused by 0.5 mM ascorbate in WT, *wei8* and the auxin import-deficient mutant *aux1* (Fig. 6C). Relative growth of *aux1* roots in the presence of ascorbate was comparable to that of *wei8* despite the presence of a fully functional auxin biosynthesis pathway in *aux1* (Fig. 6A to C). Furthermore, as expected for an auxin influx mutant, the root growth inhibition caused by ascorbate in *aux1* could not be rescued by exogenous IAA supplementation (Fig. 6B). These results highlight the importance of auxin biosynthesis and transport as a coping mechanism against the physiological impact of increased ascorbate concentration.

## Discussion

In this study, we used previously characterized strong (*vtc2*) and weak (*vtc4*) ascorbate-deficient mutants, WT, and a newly generated high-ascorbate transgenic line in which the uORF-less *VTC2* CDS is constitutively expressed, to examine the effect of altered concentrations of ascorbate on the Arabidopsis transcriptome. Ascorbate status had little effect on transcripts associated with the ascorbate biosynthesis *via* the mannose-L-galactose pathway or on ascorbate recycling enzymes. Similarly, increasing ascorbate by feeding ascorbate or its precursor L-galactono-1,4-lactone did not influence expression of ascorbate-related genes (Bulley et al., 2009; Gao et al., 2011).

The range of endogenous ascorbate concentrations in the lines studied in this work, in combination with their genome-wide transcriptomic profiles, led us to identify: (i) genes whose expression is significantly different under distinct ascorbate concentrations when compared to WT (DEGs), and (ii) genes whose expression correlates with ascorbate concentration across genotypes (PCGs and NCGs). Both *vtc2* and *vtc4* mutants showed strong reductions in the expression of *VTC2* and *VTC4* respectively and, noteworthy, we found reduced *VTC2* mRNA levels in the *vtc2/OE-VTC2* line. We hypothesize that it might be the consequence of partial transgene silencing, either due to its co-suppression in the presence of the *vtc2* mutant T-DNA (SAIL_769_H05) (Daxinger et al., 2008; Mlotshwa et al., 2010), or due to a transcriptional or post-transcriptional downregulation of the transgene (e.g., via reduced transcription or enhanced mRNA degradation) by the plant in an attempt to maintain ascorbate homeostasis.

The *vtc2* transcriptome was the most dissimilar to that of WT among all lines studied (Fig. 2A and C), with the majority of DEGs being unique to *vtc2*, while *vtc4* and *vtc2/OE-VTC2* shared over 60% of their DEGs. Surprisingly, only 37% of DEGs identified in our study for *vtc2* (*vtc2-4*) were also differentially expressed in a previous report that used the EMS allele, *vtc2-1* (Kerchev et al., 2011) (Supplementary Fig. S14; Supplementary Material 10). Although we viewed this overlap as unexpectedly low at first, it must be considered that the profiling technology (microarray used for *vtc2-1 vs.* RNA-seq used for *vtc2-4*) and the statistical cut-off (p-value<0.05 for *vtc2-1 vs.* FDR-corrected p-value<0.05 for *vtc2-4*) may increase the transcriptome differences in addition to those due to the nature of the allele (EMS *vtc2-1 vs.* T-DNA for *vtc2-4*; with Lim et al., 2016 suggesting that *vtc2-*1 has a second cryptic mutation independent from that disrupting *VTC2* that could be responsible for the growth defects of this EMS mutant) and potential differences in the growth conditions. To reconcile such differences and find commonalities, we generated a list of *vtc2* core DEGs that comprise the most reproducible *vtc2* DEGs across both studies, which is a valuable resource for the ascorbate and redox homeostasis research community (Supplementary Material 10). Based on gene ontology analysis, we found that the *vtc2* transcriptional program exhibited signatures of repressed abiotic stress responses and enhanced defense against biotic stress.

These results were somewhat expected since it was previously reported that: (i) ascorbate is involved in plant abiotic stress response and tolerance (Veljović-Jovanović et al., 2017), and (ii) *vtc1* and *vtc2* have constitutively higher transcript levels of defense genes and increased resistance to biotrophic pathogens (Barth et al., 2004; Pavet et al., 2005; Mukherjee et al., 2010). This previous work suggested that elevated H_2_O_2_ levels in these lines due to ascorbate deficiency may act as a priming mechanism that stimulates salicylic acid accumulation and increases disease resistance (Mukherjee et al., 2010). However, it was proposed that this may not be the case in *vtc4-1*, which responds more subtly to *Pseudomonas syringae* infection (Mukherjee et al., 2010). It remains to be determined whether our *vtc2/OE-VTC2* line would be more sensitive to pathogen infection due to potentially lower H_2_O_2_ levels. Consistently, we found that a large number of *WRKY* TFs were present among *vtc2*-induced genes (Supplementary Material 3), and these proteins are known to participate in complex regulatory networks that include other *WRKYs* to respond to pathogens and oxidative stress (Javed and Gao, 2023). Interestingly, no such widespread induction of *WRKY*s was observed in the milder mutant *vtc4*.

Three of the *vtc2*-induced *WRKY*s, *WRKY46*, *WRKY70*, and *WRKY53,* have been reported to have overlapping, as well as synergistic functions to coordinate basal resistance to *P. syringae* (Hu et al., 2012). In our study, we found that *WRKY46* was within “Responding as expected” class of NCG. Based on our analysis of O’Malley and collaborators’ DAP-seq TF-DNA binding data, *WRKY46* is targeted by WRKY18, 28, and 50; *WRKY53* is targeted by more than 20 different WRKYs (including WRKY18, 28, 50, and 70), but *WRKY70* is not targeted by any of these proteins. Therefore, it is possible that different ascorbate concentrations are translated into different WRKY-mediated transcriptional programs accounting for the differences observed between *vtc2* and *vtc4*. Thus, we investigated whether WRKY-targeted PCGs and NCGs were within their respective “Responding as expected” classes. However, a large number of WRKY-targeted NCGs did not respond to ascorbate in an expected manner, suggesting that it is unlikely that WRKYs are directly responsible for adjusting gene expression levels to ascorbate concentration (Supplementary Fig. S5D; Supplementary Materials 7 and 8). Consistent with the fact that we could not identify any subset of TFs that could explain genes responding in a proportional manner to ascorbate, a custom plot combining a gene’s response to ascorbate (excess *vs.* depletion) and its sensitivity to depletion (FC in *vtc4* compared to that in *vtc2*) rendered unpopulated the sector v of the plot —where *expected response* to ascorbate and *expected sensitivity to depletion* intersect (Supplementary Fig. S8A and B). It must be considered, however, that many WRKYs regulate and are regulated by hydrogen peroxide (Chen et al., 2019; Javed and Gao, 2023). Therefore, despite ascorbate and ROS playing opposite roles controlling the redox state of the cell, the abovementioned transcriptional changes that WRKYs may regulate, and the responses observed for PCGs and NCGs, could be triggered by altered ROS concentration rather than directly by ascorbate. From these analyses, we do not have enough information to infer whether the regulatory effect that ascorbate has on gene expression relies on specific TRs or signaling cascades like that of hormone signaling pathways. However, our results highlighted the importance of WRKYs in the transcriptional response to ascorbate, and we consider them primary candidates to focus on in the future to study transcriptional reprogramming upon altered ascorbate homeostasis.

The effects of ascorbate on mRNA levels of genes involved in ascorbate biosynthesis, catabolism, recycling, or regulation were mostly negligible. However, we identified 197 genes (90 PCGs and 107 NCGs) whose response was significantly altered in *vtc2* but reverted to WT by *vtc2/OE-VTC2* (Fig. 3A to D; Supplementary Material 6A and B). Among the PCGs with the strongest relative response (44-50%), we found genes that were uncharacterized like *FAD/NAD(P)-BINDING OXIDOREDUCTASE*/*AT5G11330*. Additionally, we found that *DOG1* (Fig. 3E; Supplementary Material 6A)— a well-characterized regulator of seed dormancy (Graeber et al., 2014; Footitt et al., 2020; Krüger et al., 2025) that had not been functionally linked to ascorbate before— was correlated with endogenous ascorbate in our study (Fig. 3E; Supplementary Material 6A) and with exogenously applied ascorbate precursor L-galactono-1,4-lactone (Gao et al., 2011). In turn, *MIOX1* encodes an enzyme that was proposed to increase ascorbate biosynthesis in Arabidopsis by converting myo-inositol to D-glucuronate (Lorence et al., 2004). However, strong evidence is still required since another study was unable to reproduce this result (Endres and Tenhaken, 2009), hence casting some doubts about its relevance in ascorbate biosynthesis. On the other hand, two biotic stress-related genes *AvrRpt2-INDUCED GENE1 (AIG1)* (Reuber and Ausubel, 1996; Wang et al., 2019) and *PATHOGENESIS-RELATED GENE5 (PR5)* (Uknes et al., 1992; Thomma et al., 1998; Seo et al., 2008), along with other two uncharacterized genes, namely *AT3G15536* and *AT5G13320,* led the strongest relative responses among NCGs (64%-81%; Fig. 3F; Supplementary Material 6B). The molecular mechanisms underpinning such responses are yet to be determined, and we believe that the genes listed in Supplementary Material 6 provide a valuable group of candidates to further prospect for putative regulatory elements that may account for the correlation between ascorbate concentration and gene expression, hence opening the possibility of building synthetic ascorbate-responsive reporters.

The increasing interest in the regulation of ascorbate biosynthesis in the last few years (Conklin et al., 2013; Laing et al., 2015; Fenech et al., 2021; Aarabi et al., 2023; Bournonville et al., 2023) brings about a new and exciting research front that considers a potential interplay with hormone signaling. In this regard, our expression profiling analysis of genes correlated with ascorbate concentrations led us to find a number of auxin-related genes among PCGs identified in this work, such as *TAA1* (Stepanova et al., 2008; Tao et al., 2008; Yamada et al., 2009), *RGF9* (Whitford et al., 2012), *IAA17* (Leyser et al., 1996; Kubalová et al., 2024), *TOL4* (Korbei et al., 2013) and *AIR12* (Neuteboom et al., 1999; Wang et al., 2021) (Fig. 3G; Supplementary Figs. S8A and B and S9). To validate these results, we examined the effect of exogenous ascorbate on the expression of the key auxin biosynthetic gene *TAA1* using a complemented Arabidopsis *wei8* mutant expressing a recombineering construct, *TAA1p:YPet-gTAA1*. We observed a concomitant increase of *YPet-TAA1* and *DR5* activity in response to ascorbate concentration in a light-dependent manner (Fig. 4A and D). This observation aligns well with the finding that in our transcriptomic dataset, *TAA1* expression was positively correlated with ascorbate concentration (Fig. 3C; Supplementary Figs. S8A and S9A). These results are also consistent with a previous report showing that the growth media supplementation with ascorbate or its precursor L-galactose led to a slight increase in auxin levels in Arabidopsis (Bulley et al., 2021). However, we noticed a mild reduction of root growth in *wei8* mutant in response to ascorbate not only in light-but also in dark-grown plants (Fig. 4G and H). These results may seem counterintuitive, considering that the ascorbate effect on *TAA1* expression was light-dependent (Fig. 4A to D), and ascorbate did not affect root length in WT seedlings grown in the dark. These observations are nevertheless compatible with a model in which both light and basal levels of auxin play important roles in modulating the effects of ascorbate on root length. Interestingly, we found that even in the absence of *TAA1*-encoded tryptophan aminotransferase activity, ascorbate treatment triggered an increase in auxin response, suggesting the possible involvement of other *TAA1* gene family members (Fig. 4B). In fact, we observed that the *TAA1* paralog, *TAR2*, but not *TAR1*, displayed a light-dependent increase in expression upon exogenous ascorbate supplementation (Fig. 5A and B), and the effect of ascorbate on root growth of two different allelic versions of *tar1 tar2* was intermediate between that of WT and *wei8* (Fig. 5C and D), confirming that *TAR2* is likely to be responsible for the ascorbate-mediated *DR5* induction in *wei8*.

Consistent with the idea that auxin is required to cope with excessive ascorbate, exogenous IAA supplementation of WT and *wei8* mutant alleviated their root growth defects in ascorbate-supplemented plates, whereas the roots of an auxin import mutant, *aux1,* were not complemented by IAA (Fig. 6B). However, since IAA did not fully restore the root growth inhibition triggered by ascorbate in *wei8* roots in ascorbate, we hypothesized that exogenous auxin taken up by *wei8* roots is not properly channeled towards the cell types where auxin is needed to cope with an increase in ascorbate. The observation that ascorbate-triggered root growth inhibition is similar in *wei8* and the auxin transport mutant *aux1*, and stronger than that in WT, suggests that auxin transport, in addition to local auxin production, is required for a normal response to ascorbate. In other words, auxin source cells (where local biosynthesis occurs through TAA1 and TAR2 activities in response to high ascorbate concentrations) and the cells that require auxin to respond to increased ascorbate levels are likely not the same. Instead, the root requires a functional AUX1 to deliver IAA from *TAA1/TAR2*-expressing cells into auxin sink cells. These results are consistent with previous reports that indicate that a combination of local auxin biosynthesis and transport is important to create morphogenic auxin gradients (Brumos et al., 2018), and in our case, to ensure a proper response to an increase in ascorbate concentration. Therefore, alterations in the local auxin production or auxin influx result in an enhanced ascorbate root inhibition response (Fig. 6). Further work will be required to pinpoint the exact cells that require auxin to cope with excessive ascorbate.

These results, along with the identification of *TAA1* and its paralog *TAR2* as ascorbate-regulated auxin biosynthesis genes, enabled us to identify a boost in local auxin production as an exciting new player in the response to ascorbate in roots of light-grown Arabidopsis seedlings. Furthermore, the auxin influx carrier *AUX1* was found to be required for the plant’s ability to maintain its root growth under ascorbate excess, implicating auxin transport in the ascorbate response. We believe that this novel connection to auxin and the identification of additional ascorbate-correlated genes will fuel future studies that will not only uncover how ascorbate concentration and auxin signaling are interwoven but also illuminate the impact of ascorbate on other biological processes in light-grown plants.

## Materials and methods

### Plant Materials

All Arabidopsis (*Arabidopsis thaliana*) lines used in this study are of Col-0 ecotype (WT). Arabidopsis lines employed in this work that have been described previously include: *vtc2/ GPPp:GPP-GFP L13,* herein referred to as *VTC2p:VTC2_CDS_-GFP* (Fenech et al., 2021); *vtc2-4,* herein referred to as *vtc2* (SAIL_769_H05; Lim et al., 2016); *vtc4-4*, herein referred to as *vtc4* (SALK_077222; Torabinejad et al., 2009); *wei8-1, wei8-2, tar1-1, tar2-1* and *tar2-2* mutants and their higher order combinations (Stepanova et al., 2008) and *aux1-7* mutant was previously described and characterized (Bennett et al., 1996; Marchant et al., 1999; Marchant et al., 2002). The generation of transgenic lines *vtc2/OE-VTC2* L15 and L16 for this work is described below. Recombineering line *wei8-2/TAA1p:YPet-gTAA1* was generated after backcrossing it to WT and segregating *tar2-1* out from the original line (ABRC stock #CS72246; Brumos et al., 2018; Brumos et al., 2020b) in this work. Auxin-inducible synthetic reporter *DR5:GFP* in WT and *wei8* backgrounds (Brumos et al., 2018), and *GUS* recombineering lines for *TAR* and *YUC* auxin biosynthesis genes (Brumos et al., 2020) were also previously published.

### Plasmid construction and generation of transgenic plants

The coding sequence (CDS) of *VTC2* (without stop codon) was PCR amplified from Col-0 cDNA using Phusion High-Fidelity DNA Polymerase (New England Biolabs) and the attB1-VTC2-F (5’-GGGGACAAGTTTGTACAAAAAAGCAGGCTATGTTGAAAATCAAAAGAGTTCC-3’) and attB2-VTC2-R (5’-GGGGACCACTTTGTACAAGAAAGCTGGGTGTTCTGAAGGACAAGGCACT-3’) primers. The resulting PCR product was recombined by the BP reaction into the pDNOR221 vector (Invitrogen) following manufacturer’s instructions. The *VTC2* CDS was then subcloned by LR reaction in the gateway pGWB5 vector (Nakagawa et al., 2007) to produce the *35S:VTC2-GFP* construct. The fidelity of the construct was confirmed by sequencing. Arabidopsis *vtc2-4* mutant (SAIL_769_H05; Lim et al., 2016) was transformed using the GV3101 strain of *Agrobacterium tumefaciens* using the floral dip protocol (Clough and Bent, 1998). Transgenic lines were screened for 3:1 segregation on half-strength Murashige-Skoog (M5524 Sigma-Aldrich) plates containing 20 µg/mL Hygromycin (Invitrogen) and propagated to homozygous T3 lines. 16 independent lines were screened and two lines showing high ascorbate content were selected.

### Growth conditions

For RNA-seq analysis, Arabidopsis seeds were surface sterilized using chlorine gas by pouring 3 mL of 37% HCl into 100 mL of commercial bleach in a flask and incubated in an airtight sealed container for 4 hours. Then, seeds were aired for at least 4 hours in a laminar flow cabinet and stored at 4°C for a three-day stratification. Seeds were sown on horizontal half-strength MS (M524 PhytoTech Labs, USA) agar (0.6% [w/v]) (A0949 PanReac Applichem ITW Reagents), pH 5.7 plates supplemented with 1.5% [w/v] sucrose under sterile conditions and grown with a long-day photoperiod (16-h light/8-h darkness cycle, 22±1ᵒC, 150±50 μmol photons m^-2^ s^-1^) for 7 days. Seedlings were then transferred to soil (4:1 (v/v) soil:vermiculite) in a semi-randomized order. Rosettes of four-week-old plants grown under short-day photoperiod (8-h light/16-h darkness cycle, 22±1ᵒC, 300±70 μmol photons m^-2^ s^-1^) were collected 30 minutes after the lights were turned on and flash-frozen in liquid nitrogen. RNA from three independent replicates per genotype was extracted following the protocol described below.

For ascorbate- and auxin-related experiments, Arabidopsis seeds were surface-sterilized using a sterilization solution (50% commercial bleach (Pure Bright germicidal ultra bleach), 0.02% (v/v) Triton X-100 (Pharmacia Biotech code no. 17-1315-01)) for 5 minutes and washed four times with sterile deionized water. Seeds were stratified for three days at 4°C, sown after resuspending them in low-melting point agarose (0.7%) in water on horizontal full-strength Murashige-Skoog (MS) (Caisson Labs ref. MSP01-100LT) agar (0.6% [w/v]) (Difco agar granulated ref. 214510), pH6 supplemented with 1% [w/v] sucrose, and kept for two hours at room temperature to induce germination. The pH of ascorbate-supplemented medium was adjusted to 6 using extra empirically determined 1M KOH to accommodate 0.5mM ascorbate, and equivalent volume of 1M KCl was added to control medium. This medium was autoclaved and cooled down at room temperature before adding filter-sterilized ascorbic acid (Sigma-Aldrich ref. A5960) using 0.22µM filters for aqueous solutions (Millex-GV, Polyvinylidene fluoride (PVDF) ref. SLGV013SL) or indole-3-acetic acid (Sigma ref. I-2886) using dimethylsulfoxide (DMSO)-compatible filters (Choice Polypropylene Syringe Filters ref. CH2225). Plates with seeds for the dark experiment were then transferred to a dark chamber at 22°C and incubated for 72 hours. Plates with seeds for the light experiment were transferred to a light walk-in chamber and incubated under continuous LED light (70-100 μmol photons m^-2^ s^-1^; 2x 6000K Kihung T8 LED integrated fixture 40W + 1x FULL SPECTRUM Monios-L LED grow light full spectrum 60W) for 5 days.

### Experimental procedures

Ascorbate concentration was determined using an ascorbate oxidase activity assay as previously described (Fenech et al., 2021). Confocal imaging was performed using Carl Zeiss LSM710 (excitation: argon laser 488nm; detection: 499nm-562nm) and brightness/contrast was adjusted using FIJI. For GUS histochemical staining of auxin biosynthesis recombineering lines and *DR5*, seedlings were fixed in 90% (v/v) acetone and stained for 48 hours as previously described (Stepanova et al., 2005). Seedling images were acquired using a Q-IMAGING MicroPublisher 5.0 RTV camera coupled to a LEICA MZ125 stereo microscope, and GUS-stained seedlings were imaged using a Diagnostic Instruments Spot Insight 4 14.2 color mosaic Camera coupled to a Carl Zeiss Axioplan microscope. Seedling images and organ size measurements were acquired using an Epson Perfection V600 Photo scanner and analyzed with FIJI/ImageJ.

### RNA extraction

RNA from three independent replicates per genotype, where each biological replicate was composed of 8 randomized rosettes, was extracted with Trizol following the manufacturer’s protocol. RNA quality and integrity were validated using the Bioanalyzer 2100 (Agilent Technologies Santa Clara, CA, USA), with the RNA integrity number (RIN) of >8.0 for all biological replicates.

### RNA-seq differential expression and bioinformatics analysis

Paired-end Illumina mRNA libraries were generated using the TruSeq stranded mRNA kit according to the manufacturer’s instructions (Illumina Inc., San Diego, CA, USA) for the mutant and control lines. Libraries were sequenced in an Illumina NextSeq550 platform, and 2 x 75 bp paired-end reads were generated.

The reads were quality-filtered and trimmed using Trimmomatic version 0.36 (Bolger et al., 2014) with paired-end mode options: -threads 8 -phred33 ILLUMINACLIP:TruSeq3-PE.fa:2:30:10:2:True LEADING:3 TRAILING:3 MINLEN:36. The resulting reads were then aligned to the TAIR10 version of the *Arabidopsis thaliana* genome sequence (https://www.arabidopsis.org/) using Hisat2 version 2.1.0 (Kim et al., 2015). These read alignments (in BAM format) were used for transcript quantification with the Cuffdiff program of the Cufflinks version 2.2.1 package (Trapnell et al., 2013). The resulting read alignments were visualized and clustered using Tablet software (Milne et al., 2013) and CummeRbund R package version 2.23.0 (Goff et al., 2014). To determine DEGs, a q-value<0.05 cutoff was set. For DEGs analysis, genes identified as ID ENSRNA were filtered out.

### Principal Component Analysis (PCA) and volcano plot of RNA-seq datasets

To explore the relationship between fold-change and significance (q-value<0.05), Volcano plots were generated using the CummeRbund R package version 2.23.0 (Trapnell et al., 2012). PCA was made using a customized pipeline in RStudio using ggplot2 (Wickham et al., 2025a) and ggfortify (Horikoshi et al., 2024) R packages.

### Gene ontology (GO) categories analysis

The PANTHER database (http://go.pantherdb.org/tools/compareToRefList.jsp) was used to obtain the GO biological processes overrepresented in the input gene list (DEGs of a given mutant), using as reference list all *Arabidopsis thaliana* genes annotated in the PANTHER database (Mi et al., 2019; Thomas et al., 2022). The Fisher’s exact test with the FDR correction as calculated by the Benjamini-Hochberg procedure was used and only results with FDR<0.05 were selected for further analysis.

### Venn diagram analysis and statistical analysis of the overlap

The Venn diagram analysis was performed using “InteractiVenn” and on-line interactive Venn diagram tool and introducing the list of induced and repressed genes for each mutant as input (http://www.interactivenn.net/index.html<u>#</u>) (Heberle et al., 2015).

The statistical significance of the overlap between two groups of genes was calculated using an online tool (http://nemates.org/MA/progs/overlap_stats.html). The complete description of how the enrichment (representation factor) and the associated probability (p-value) were calculated can be found in this web page (http://nemates.org/MA/progs/representation.stats.html).

### Correlation analysis

We filtered out genes with Fragments Per Kilobase of transcript per Million mapped reads (FPKM)<1 in at least one of their replicates, and we kept the genes whose average FPKM values followed the desired pattern of *vtc2*>*vtc4*>WT>*vtc2/OE-VTC2* to find NCGs, or *vtc2*<*vtc4*<WT<*vtc2/OE-VTC2* to find PCGs. To do that, we wrote a pipeline in RStudio utilizing dplyr (Wickham et al., 2023), R.utils (Bengtsson, 2025), and stats R packages (R Core Team, 2025). We performed Pearson’s correlation analysis with no transformation to extract genes whose response to ascorbate is linear (FPKM=±m*[ascorbate]+n). Additionally, we performed the same analysis after transforming FPKM values into Napierian logarithm (ln(FPKM)= =±m*[ascorbate]+n) for exponential response, and in the case of NCGs, we also transformed ascorbate concentration into 1/[ascorbate] to identify genes whose expression is inversely proportional to ascorbate concentration (FPKM value positively correlates with 1/[ascorbate]). Best fit for each gene is shown in Supplementary Material 5.

### Hierarchically clustered TF-target heatmap

This analysis was made using a customized pipeline in RStudio. First, we downloaded the TF-target interactions dataset published by O’Malley et al., 2016. Then, we processed transcription factor (TF) binding data, utilizing the dplyr package (Wickham et al., 2023) for data manipulation, in naïve/methylated (labelled as “col”) and unmethylated (labelled as “colamp”) DNA to create a TF-target gene database. Finally, we constructed a binary interaction matrix of TF-target pairs for hierarchical clustering using the methylated database. To do this, a matrix was initialized where rows represent target genes and columns represent TFs, with each cell indicating whether a TF binds to a target gene. The matrix was populated with 1 if a TF binds to a target gene and 0 if no binding is detected. To visualize these interactions, the binary matrix was converted into a heatmap using the ComplexHeatmap package in R (Gu et al., 2016; Gu, 2022), with a color gradient distinguishing binding (dark color) from non-binding (light color). Hierarchical clustering was applied to group TFs, while target genes were arranged based on their *vtc2* expression fold-change.

### Development of a custom plot to visualize changes in gene expression in response to ascorbate concentration

We hypothesized that, if the expression of an ideal gene (measured as FPKM) was directly proportional to the concentration of ascorbate responding as 1:1, its expression across genotypes would follow two different models: a linear model for a PCG (FPKM_genotype_ = m * [ascorbate]_genotype_, with 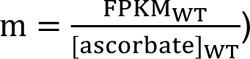, and an inverse model for a 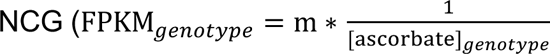, with m = [ascorbate]_WT_ * FPKM_WT_). Using this, we predicted that in the *vtc2* mutant (20% of WT ascorbate), this PCG’s expression would be 0.2 times that of WT, while in *vtc2/OE-VTC2* (165% of WT ascorbate), it would be 1.65 times higher. Conversely, an ideal NCG would show 5-fold higher expression in *vtc2* and 0.61-fold in *vtc2/OE-VTC2* (inverse values). In log2 scale, genes with fold changes of (−2.32, 0.72) for PCGs or (2.32, −0.72) for NCGs would be considered correlated 1:1. The red line in Supplementary Fig. S5A and B represents all theoretically possible genes responding to ascorbate concentration more weakly than a 1:1, but still keeping a balanced proportion between *vtc2* and *vtc2/OE-VTC2*, and genes falling on this line are considered to respond to ascorbate as expected. To experimentally identify these genes, we established a confidence interval based on the highest and lowest observed ascorbate fold changes in *vtc2* and *vtc2/OE-VTC2*, assuming that the true mean could be any of the three measured values per genotype. This confidence range, determined by comparing extreme ascorbate values across replicates, defined two boundary points, and the lines connecting (0,0) to these points represent the upper and lower empirical response limits. The same rationale was utilized to build the plots displayed in Supplementary Fig. S7A and B. Then, both plots were combined in Supplementary Fig. S8A and B, by representing the slope values that each gene had in Supplementary Fig. S5A and B (represented in y-axis) and Supplementary Fig. S7A and B (represented in x-axis). All visual representations of gene expression in response to ascorbate concentration were performed in RStudio utilizing dplyr (Wickham et al., 2023), ggplot2 (Wickham et al., 2025a), ggpubr (Kassambara, 2023a), svglite (Wickham et al., 2025b), and viridisLite (Garnier et al., 2024) R packages.

### Statistical analysis

We made different *ad hoc* pipeline scripts in R to perform statistical analyses utilizing car (Fox and Weisberg, 2019), pwr (Champely et al., 2020), rstatix (Kassambara, 2023b), dunn.test (Dinno, 2024), and multcompView (Graves and Dorai-Raj, 2024) packages. Firstly, we tested whether the dataset was normally distributed (Shapiro-Wilk test) and homoscedastic (Levene test) prior to running the hypothesis contrast analyses. If a dataset was homoscedastic but had strong statistical power (power=1 – β > 0.8, with β being the probability of accepting the null hypothesis when it is actually false), we performed ANOVA followed by post-hoc Tukey test for α=0.05. If not meeting the previously stated requirements, we transformed the data using log_10_ and retested for homoscedasticity and power. If after data transformation the requirements were not met, we performed non-parametric Kruskal-Wallis test, followed by Dunn’s test for α=0.05.

## Acknowledgements

We thank Dr. Deyu Xie (NC State University, USA) for generously sharing an aliquot of ascorbic acid with us to run the experimental validation of this work. Generative Artificial Intelligence (Open AI) was utilized to assist in the troubleshooting of writing the RStudio scripts utilized to analyze statistically and plot the results.

## Author contributions

M.F., V.A.-S., M.A.B., N.S., ANS, and JMA designed the research. D.A. generated the *vtc2/OE-VTC2* line in N.S.’s laboratory, M.F. designed and performed the experiments, C.M-P. extracted RNA for RNA-seq, assembled the transcriptome, and analyzed the data. M.F., V.Z., J.M.-C., and I.M. curated and analyzed the data. M.F. and V.Z. generated and curated the R scripts necessary to run correlation analyses and for the visualization of the results. M.F. and V.A.-S. wrote the manuscript with the contribution of all other authors.

## Data availability

The data generated in this publication have been deposited in NCBI’s Gene Expression Omnibus (Edgar et al., 2002) and are accessible through GEO Series accession number GSE296827 (https://www.ncbi.nlm.nih.gov/geo/query/acc.cgi?acc=GSE296827). All R scripts utilized to analyze and plot correlation and postcorrelation analyses are available in GitHub (doi: 10.5281/zenodo.15424810).

---

**Supplementary Figure S1.**
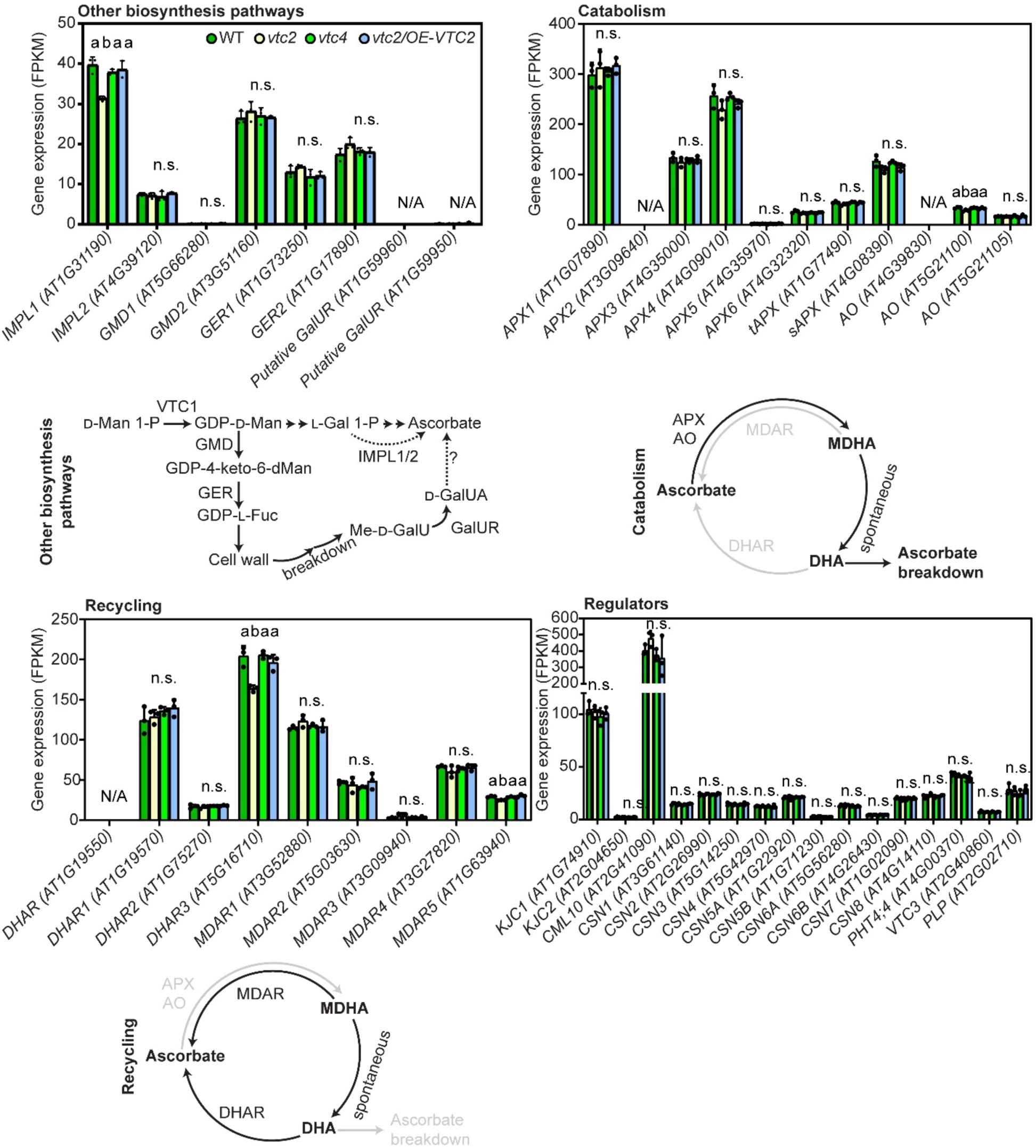
Expression of genes related to ascorbate homeostasis. Individual gene plots are available in Supplementary Fig. S2. Different letters denote statistically significant differences (One-Way ANOVA, Tukey post-hoc test, α=0.05). Distributions were tested to meet normality (Shapiro–Wilk’s test) and homoscedasticity (Levene’s test) prior to ANOVA. If not meeting any of these requirements, data was transformed using logarithm and, if meeting the requirements, proceeded to perform ANOVA. Otherwise, Kruskal-Wallis followed by Dunn’s post-hoc test were performed. *IMPL1: MYO-INOSITOL MONOPHOSPHATASE LIKE 1. IMPL2/HISN7: MYO-INOSITOL MONOPHOSPHATASE LIKE 2/HISTIDINE BIOSYNTHESIS 7, GMD1: GDP-D-MANNOSE-4,6-DEHYDRATASE 1; GER1: GDP-4-KETO-6-DEOXYMANNOSE-3,5-EPIMERASE-4-REDUCTASE, GalUR: D-GALACTURONATE ACID REDUCTASE. APX: ASCORBATE PEROXIDASE (tAPX: tylakoidal APX, sAPX: stromatic APX), AO: ASCORBATE OXIDASE, MDHAR, MONODEHYDROASCORBATE REDUCTASE; DHAR, DEHYDROASCORBATE REDUCTASE, KJC1: KONJAC1, CML10: CALMODULIN-LIKE10, CSN1: COP9 SIGNALOSOME COMPLEX SUBUNIT 1, PHT4.4: INORGANIC PHOSPHATE TRANSPORTER, VTC3: DUAL FUNCTION PROTEIN KINASE::PROTEIN PHOSPHATASE.* FPKM: Fragments Per Kilobase of transcript per Million mapped reads

**Supplementary Figure S2.**
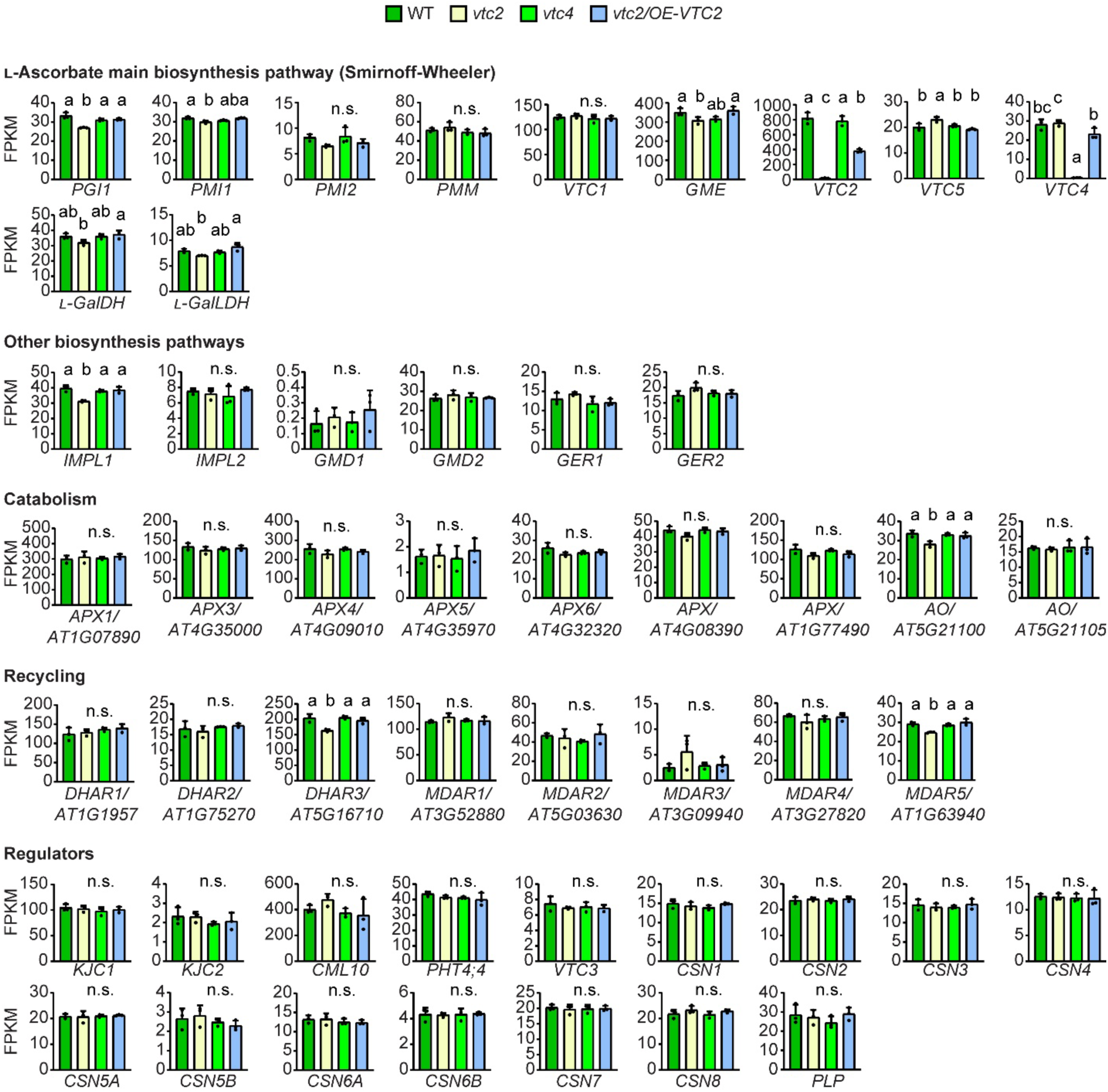
Expression of individual genes related to ascorbate homeostasis. Different letters denote statistically significant differences (One-Way ANOVA, Tukey post-hoc test, α=0.05). Distributions were tested to meet normality (Shapiro–Wilk’s test) and homoscedasticity (Levene’s test) prior to ANOVA. If not meeting any of these requirements, data was transformed using logarithm and, if meeting the requirements, proceeded to perform ANOVA. Otherwise, Kruskal-Wallis followed by Dunn’s post-hoc test were performed. *PMI: PHOSPHOMANNOSE ISOMERASE, PMM: PHOSPHOMANNOMUTASE, VTC1: VITAMIN C DEFICIENT1 (GDP-D-MANNOSE PYROPHOSPHORYLASE), GME: GDP-D-MANNOSE-3′,5′-ISOMERASE, VTC2: GDP-L-GALACTOSE PHOSPHORYLASE, VTC4: L-GALACTOSE-1-PHOSPHATE PHOSPHATASE, L-GalDH: L-GALACTOSE DEHYDROGENASE, L-GalLDH: L-GALACTONO-1,4-LACTONE DEHYDROGENASE. IMPL1: MYO-INOSITOL MONOPHOSPHATASE LIKE 1. IMPL2/HISN7: MYO-INOSITOL MONOPHOSPHATASE LIKE 2/HISTIDINE BIOSYNTHESIS 7, GMD1: GDP-D-MANNOSE-4,6-DEHYDRATASE 1; GER1: GDP-4-KETO-6-DEOXYMANNOSE-3,5-EPIMERASE-4-REDUCTASE. APX: ASCORBATE PEROXIDASE, AO: ASCORBATE OXIDASE, MDHAR, MONODEHYDROASCORBATE REDUCTASE; DHAR, DEHYDROASCORBATE REDUCTASE, KJC1: KONJAC1, CML10: CALMODULIN-LIKE10, PHT4.4: INORGANIC PHOSPHATE TRANSPORTER, VTC3: DUAL FUNCTION PROTEIN KINASE::PROTEIN PHOSPHATASE, CSN1: COP9 SIGNALOSOME COMPLEX SUBUNIT 1, PLP: PAS/LOV (Per-ARNT-Sim/Light-Oxygen-Voltage) PROTEIN.* FPKM: Fragments Per Kilobase of transcript per Million mapped reads

**Supplementary Figure S3.**
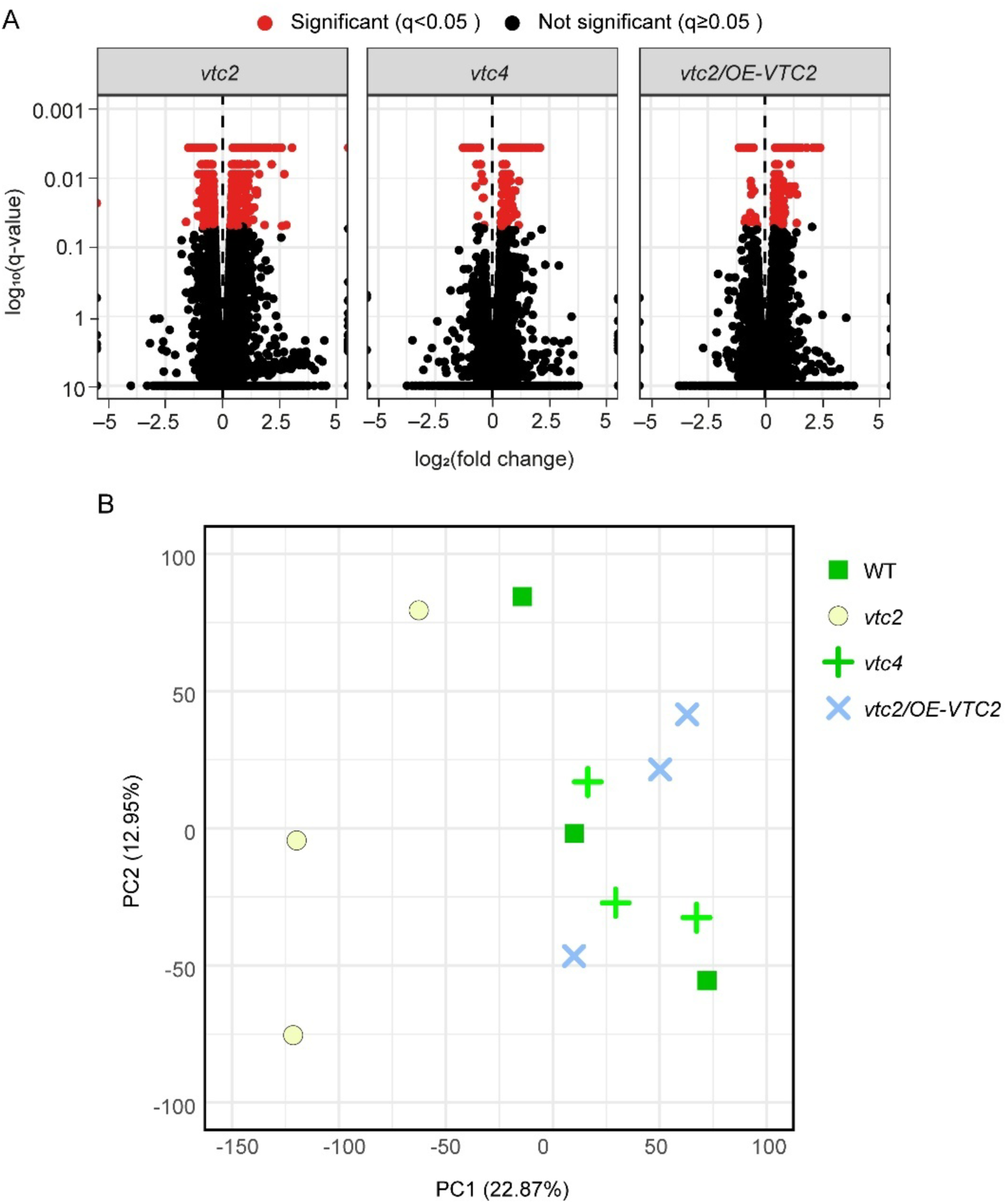
Transcriptome profiles of the ascorbate-altered Arabidopsis lines. A) Volcano plots showing that the statistically significant DEGs (red) are more abundant in *vtc2* compared to *vtc4* or *vtc2/OE-VTC2*. Fold change between each ascorbate mutant and WT is presented in log2 scale, whereas FDR-corrected p-values (q-values) are presented in log10 scale. Significance cutoff was set at α=0.05. Among Differentially expressed genes (DEGs) (q-value<0.05, red dots) across genotypes, the smallest log2(fold-change)=0.33 (1,25-fold) for the induced genes and −0.33 (0.8-fold) for repressed genes. B) Principal Component Analysis plot displaying all replicates for each of the four genotypes.

**Supplementary Figure S4.**
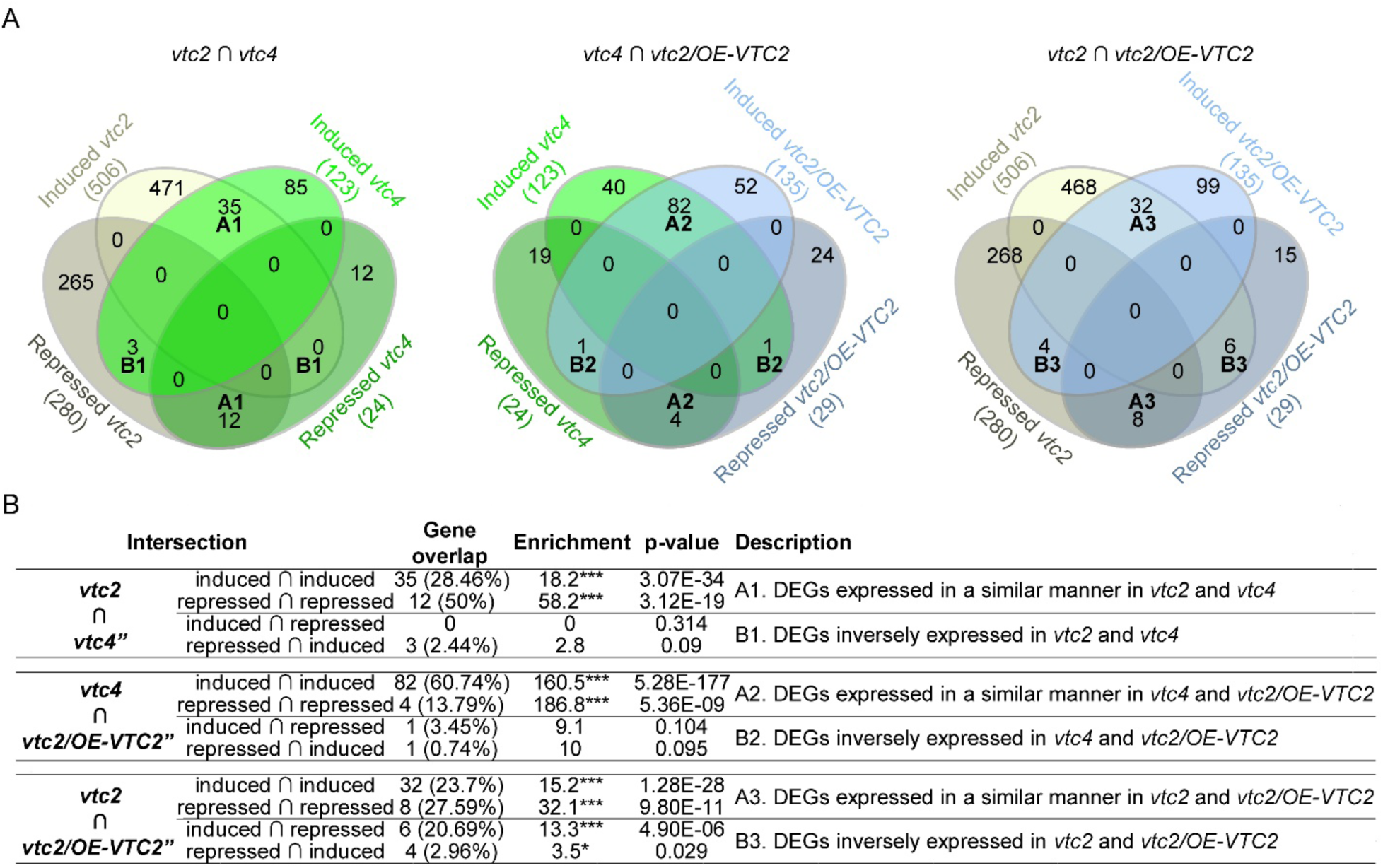
A) Venn diagrams depicting the pairwise overlap of DEGs between the three ascorbate-altered lines. A) Absolute number of genes overlapping between two plant lines. The ∩ symbol denotes intersection. For any two sets of X and Y, X∩Y is read as X intersection with Y. Group names A1-A3 and B2-B3 are described in B). B) Statistical analysis of the DEG overlap. The percentage of the overlapped genes was calculated by dividing the number of overlapped genes by the number of genes in the genotype marked with the symbol (“) and multiplied by 100. The enrichment is calculated by dividing the number of overlapping genes by the expected number of overlapping genes (if the overlap was random) drawn from two independent groups. Therefore, values >1 denote more overlap than expected, <1 less overlap than expected and, =1 indicates the expected overlap between two independent groups of genes. Level of statistical significance of the overlap is noted as * (p-value<0.05) and *** (p-value<0.001).

**Supplementary Figure S5.**
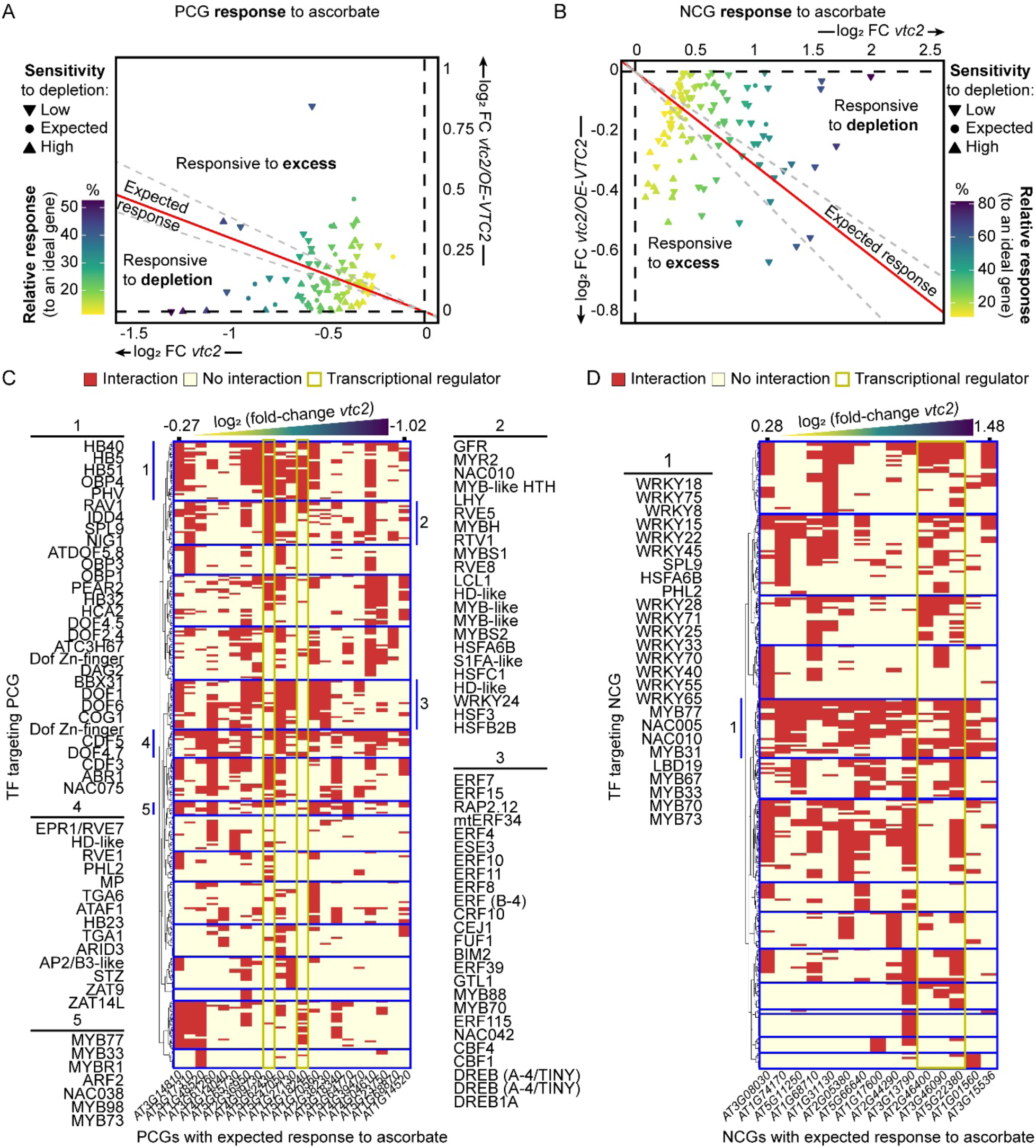
Ascorbate-correlated genes show different types of response to ascorbate concentration. A) Positively correlated genes (PCGs). B) Negatively correlated genes (NCGs). To establish the identity of the plot area, the following scenarios were considered: a theoretical gene would experience a 1:1 change in expression following a change in ascorbate concentration. Thus, this ideal PCG would be 5-fold repressed in vtc2 (20% of WT ascorbate content), and 1.65-fold induced in vtc2/OE-VTC2 (165% of WT ascorbate content), whereas an ideal NCG would be 5-fold induced in vtc2 and 1.65-fold repressed in vtc2/OE-VTC2. Taking log2, these fold changes would be −2.34 for vtc2 and 0.72 for vtc2/OE-VTC2 (PCG) or 2.34 and −0.72, respectively, as an NCG, hence defining the coordinates for an ideal PCG (−2.34, 0.72) and for an ideal NCG (2.34, −0.72). The line than links these point with the (0,0) defines the ideal proportion in the change of expression given the change in ascorbate concentration. The distance between (0,0) and (2.34, −0.72) represents the 100% of a 1:1 response (see Supplementary Fig. S6 for further details). Genes whose expression is changed in both vtc2 and vtc2/OE-VTC2 lines following this proportion (i.e., log2FC vtc2/OE-VTC2: log2FC vtc2 remains constant) are classified as correlated genes responding as expected and lay near the red line within the calculated error margins (gray dotted lines). In turn, a given gene with an expression FC vtc2 close to 0 implies that a severe ascorbate depletion does not affect its expression, and therefore, its correlation with ascorbate concentration can only be by due to the response to ascorbate excess. Following the same rationale, correlated genes with FC vtc2/OE-VTC2 close to 0 show WT levels of expression under the accumulation of ascorbate, and therefore, these genes are classified as responsive to ascorbate depletion. The symbols represent the sensitivity of a given gene to ascorbate depletion, calculated as log2FC vtc4: log2FC vtc2, graphically represented in Supplementary Fig. S6. C, D) Transcription Factors (TF) interacting with the promoters of PCGs (C) and NCGs (D) with expected response (i.e., proportional to (−2.34, 0.72) or (2.34, 0.72)), extracted from O’Malley et al., (2016) using methylated DNA as template. Detected interaction between the TF and the promoter of the gene is colored in red, whereas no interaction is colored in light yellow. Transcriptional regulators that are correlated to ascorbate concentration as expected are highlighted with a yellow rectangle. Targeted correlated genes are arranged from the mildest (left) to the strongest (right) expression fold-change in the vtc2 mutant. TF clustering distances were calculated and plotted using Ward.D2 method in ComplexHeatmap R package (Gu et al., 2016; Gu, 2022).

**Supplementary Figure S6.**
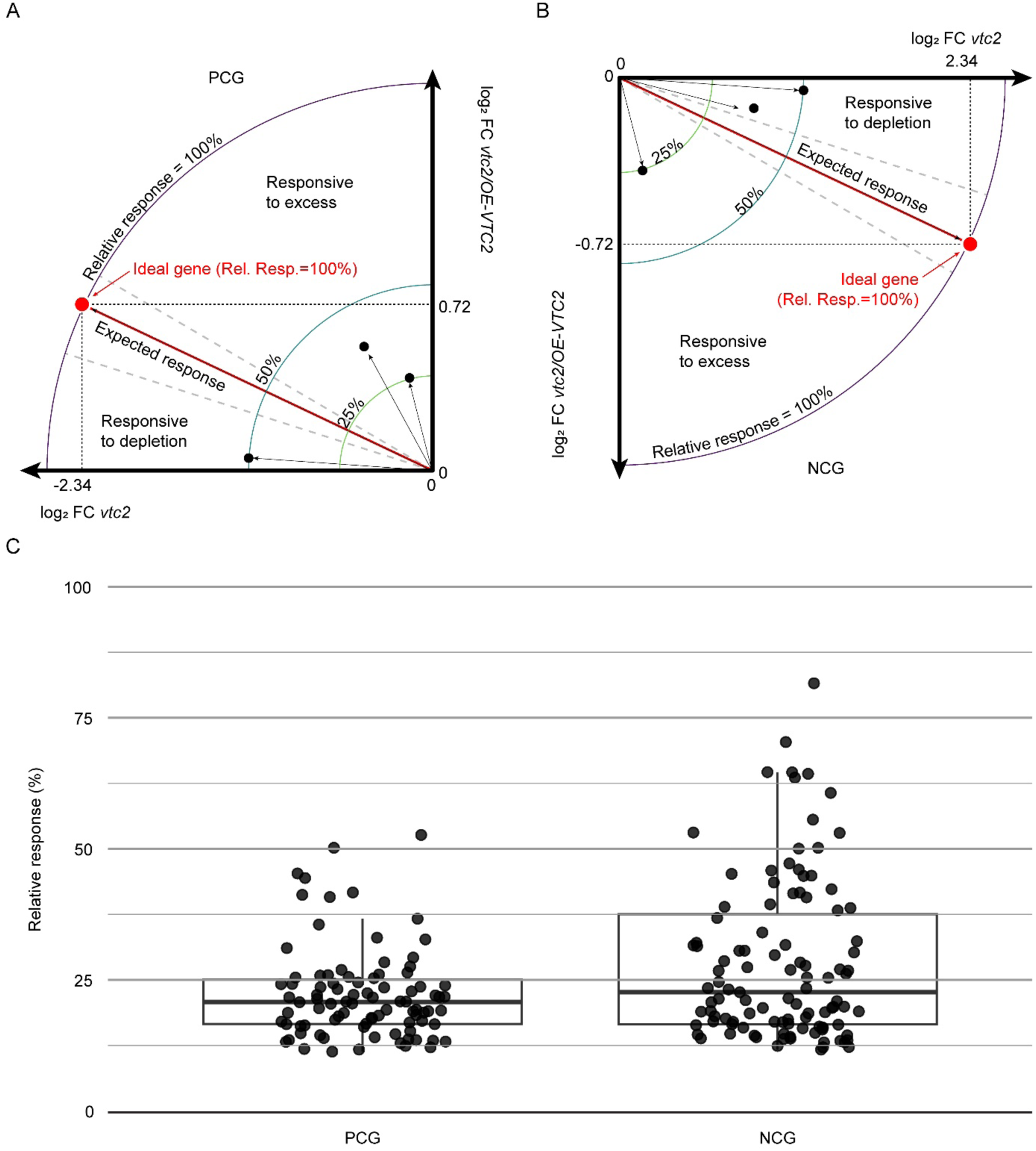
Relative response of ascorbate correlated genes compared to an ideal ascorbate-regulated gene. A theoretical ideal gene would experience a change in expression proportional to the change in ascorbate concentration. A) A Positively Correlated Gene (PCG) responding 1:1 to ascorbate concentration would experience a 5-fold repression in *vtc2* (20% of WT ascorbate content) and a 1.65-fold induction in *vtc2/OE-VTC2* (165% of WT ascorbate content), whereas a B) Negatively Correlated Gene (NCG) would be 5-fold induced in *vtc2* and 1.65-fold repressed in *vtc2/OE-VTC2*. Taking log2Fold change (FC) of *vtc2*’s and *vtc2/OE-VTC2*‘s ascorbate concentration, an ideal PCG would lay on the (−2.32, 0.72) and an ideal NCG on the (2.32, −0.72). Hence, the distance between (0,0) and these two points set the 100% of a 1:1 response of gene expression to ascorbate concentration. C) Distribution of relative responses observed in ascorbate-correlated genes. We considered the distance between the ideal gene and the (0, 0) as the 100% relative response to ascorbate concentration (also noted as Rel. Resp.) and the relative response of all correlated genes (black thin arrows) were calculated using the Pythagorean theorem and normalized to the ideal gene.

**Supplementary Figure S7.**
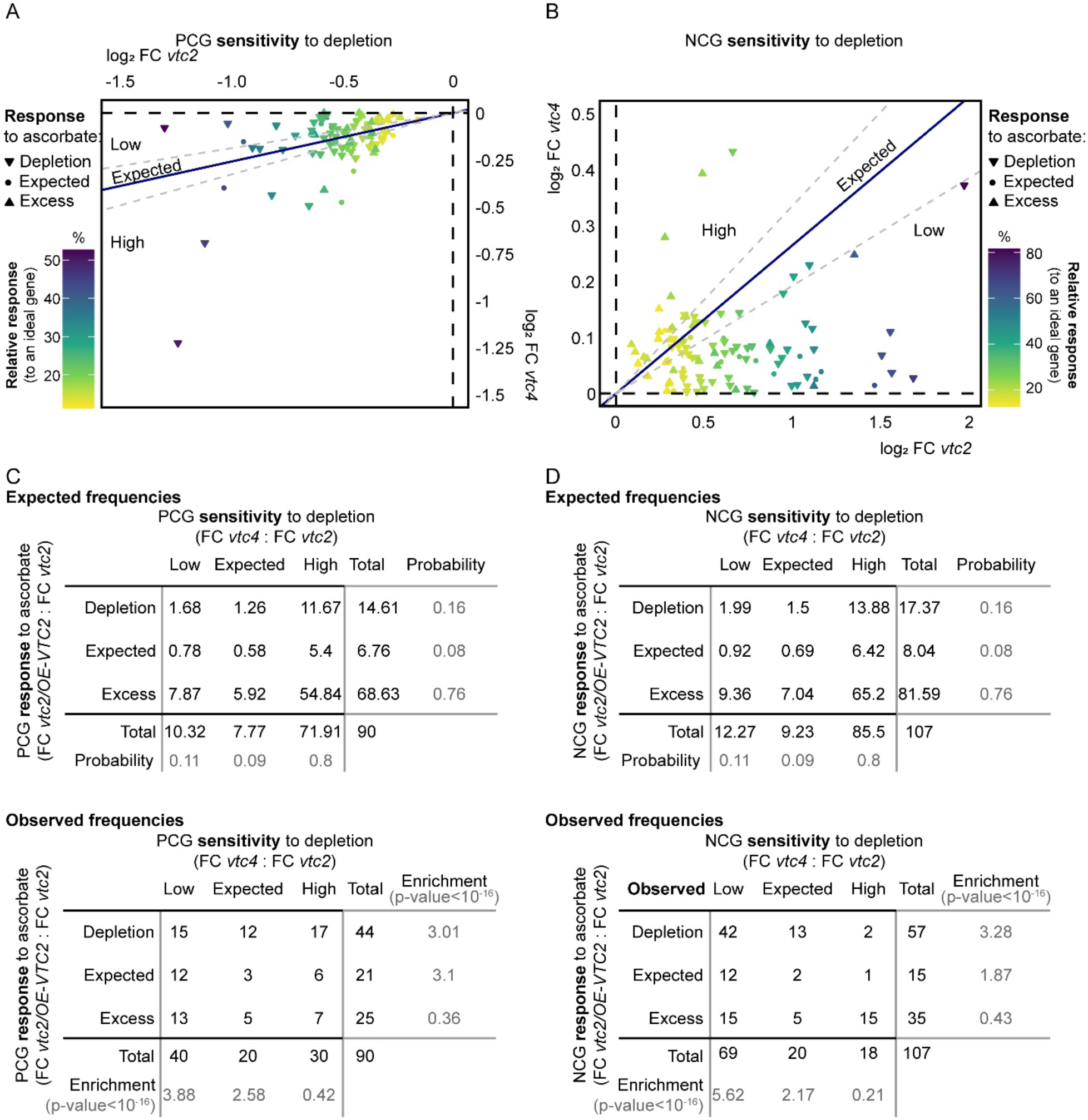
Ascorbate-correlated genes are enriched in the response to depletion. Gene expression sensitivity to ascorbate depletion of A) PCGs and B) NCGs. To establish the identity of the plot area, we considered the following premises. A theoretical ideal gene would experience a change in expression proportional to the change in ascorbate concentration. An ideal PCG would be 5-fold repressed in *vtc2* (20% of WT ascorbate content), and 1.53-fold repressed in *vtc4* (65% of WT ascorbate content), whereas an ideal NCG would be 5-fold induced in *vtc2* and 1.53-fold induced in *vtc4*. Taking log2 Fold change (FC) of *vtc2*’s and *vtc4*’s ascorbate concentration, an ideal PCG would experience a change in expression that would be represented as point (−2.32, −0.61) and an ideal NCG as (2.32, 0.61). The navy-blue line represent all values that keep the log2FC *vtc2*: log2FC *vtc4* equal to 2.32/0.61. Therefore, those genes lying near the blue line within the error margins (gray dashed lines) are considered genes with an expected sensitivity to ascorbate depletion. Genes with a greater ratio experienced a greater-than-expected change in expression under a mild ascorbate deficiency (*vtc4*), and therefore, we considered these genes had high sensitivity to ascorbate depletion. Conversely, genes whose expression FC in *vtc4* is closer to 0 would represent genes that remained unaltered under mild ascorbate deficiency, and therefore, they are classified as genes with low sensitivity to ascorbate depletion. C,D) Expected (top panel) and observed (lower panel) frequencies of PCGs (C) and NCGs (D) belonging to each category of response to ascorbate (based on log2FC *vtc2/OE-VTC2*: log2FC *vtc2*, see Supplementary Fig. S5) and sensitivity to its depletion (log2FC *vtc2*: log2FC *vtc4*), also available in Supp. Material 6A-B. The enrichment is calculated as the total number of observed genes belonging to a category divided by the total number of expected genes belonging to the same category. The expected number of genes for a given category were calculated multiplying the probability of belonging to that category by chance by the total number of PCGs (90) or NCGs (107). The probability of belonging to a given category by chance was calculated using a Monte Carlo approach based on the proportion that each category takes in the plot. To do that, a blank plot with equal-length axes containing the “expected” area was printed and proportions were estimated based on weight using a precision scale. To guarantee accuracy in the proportions, a circular line whose radius equals the length of the axes was inscribed to ensure that probabilities estimated consider genes with the same relative response.

**Supplementary Figure S8.**
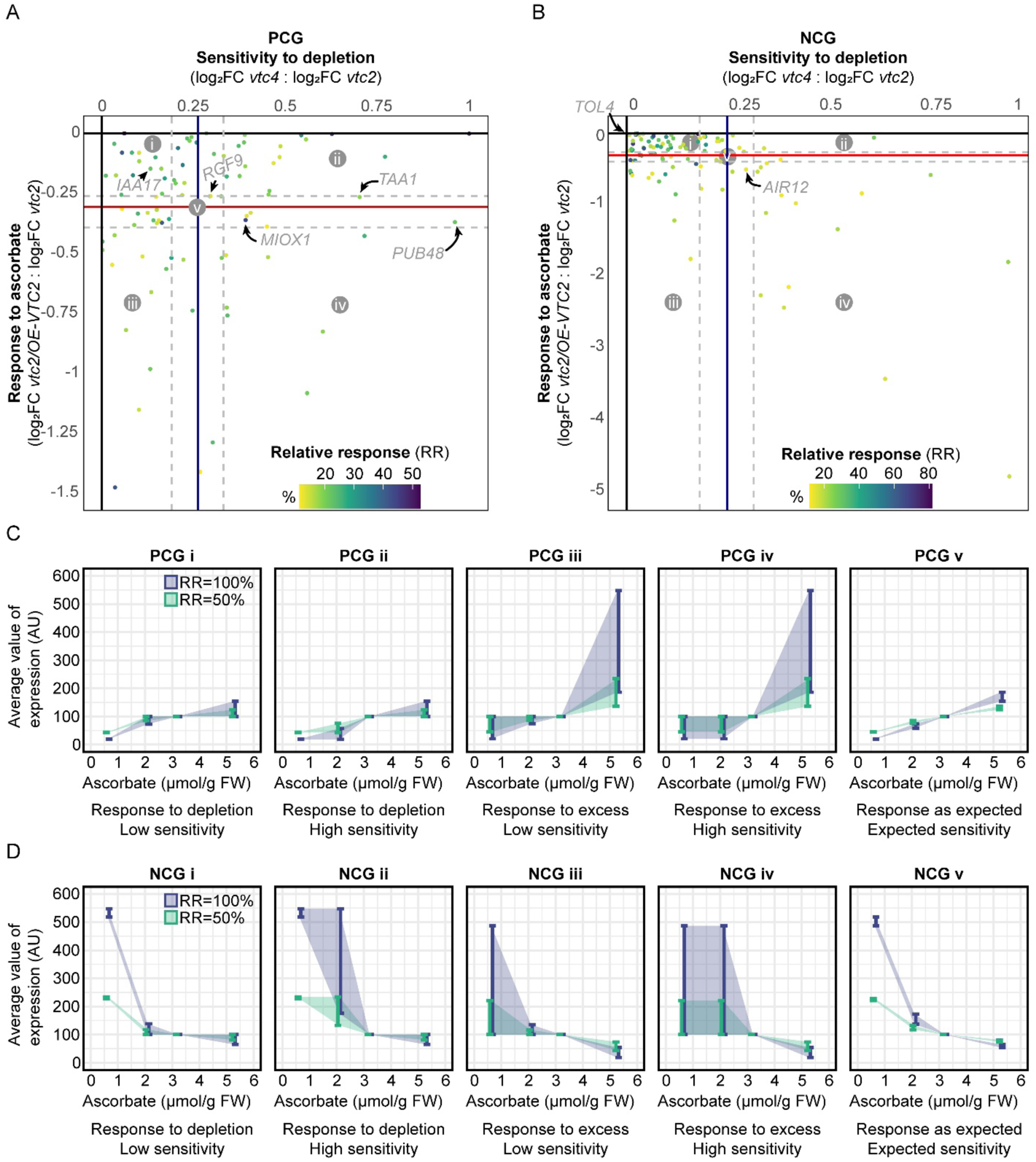
The expression patterns of ascorbate-correlated genes. A) Positively Correlated Genes (PCGs). B) Negatively Correlated Genes (NCGs). The y-axis represents a gene’s response to changes in ascorbate concentration and it is obtained from Supplementary Fig. S5A and B by dividing gene expression FC in *vtc2/VTC2-GFP* by that in *vtc2*. The x-axis represents a gene’s sensitivity to ascorbate depletion, and it is obtained from Supplementary Fig. S7A and B by dividing a gene’s expression FC in *vtc4* by that in *vtc2*. To establish the identity of the plot area, the following scenarios were considered. A theoretical ideal gene would experience a change in expression proportional to the change in ascorbate concentration. Thus, the expression FC of an ideal PCG would be 0.2 in *vtc2*, 0.65 in *vtc4*, and 1.65 in *vtc2/OE-VTC2*, whereas for an ideal NCG it would be 5, 1.53, and 0.61 for *vtc2*, *vtc4*, and *vtc2/OE-VTC2*, respectively. Therefore, after taking log2, an ideal gene would show a FC *vtc2/OE-VTC2*: FC *vtc2* = −0.31 (horizontal red line) and a FC *vtc4*: FC *vtc2* = 0.26 (vertical navy-blue line). Thus, the horizontal red line intersecting the y-axis and the dashed gray horizontal lines represent the expected response and the error margins of calculation, respectively, like in Supplementary Fig. S5A and B, and the vertical navy-blue line represents the expected sensitivity of a gene and it is homolog to that in Supplementary Fig. S7C and D) Calculated interval where the average expression (arbitrary units, AU) for a given genotype can be compared to WT, based on the sector they lay in as a PCG (C) or as a NCG (D), considering two relative responses (RR, defined as the fold-change in expression: fold-change in ascorbate concentration compared to a 1:1 ratio). IAA17: *INDOLE-3-ACETIC ACID INDUCIBLE 17 (AT1G04250), RGF9: ROOT MERISTEM GROWTH FACTOR 9 (AT5G64770), MIOX1: MYO-INOSITOL OXYGENASE 1 (AT1G14520), TAA1: TRYPTOPHAN AMINOTRANSFERASE OF ARABIDOPSIS 1 (AT1G70560), PUB48: PLANT U-BOX 48 (AT5G18340), TOL4: TOM1-LIKE 4 (AT1G76970), AIR12: AUXIN-INDUCED IN ROOT CULTURES 12 (AT3G07390)*.

**Supplementary Figure S9.**
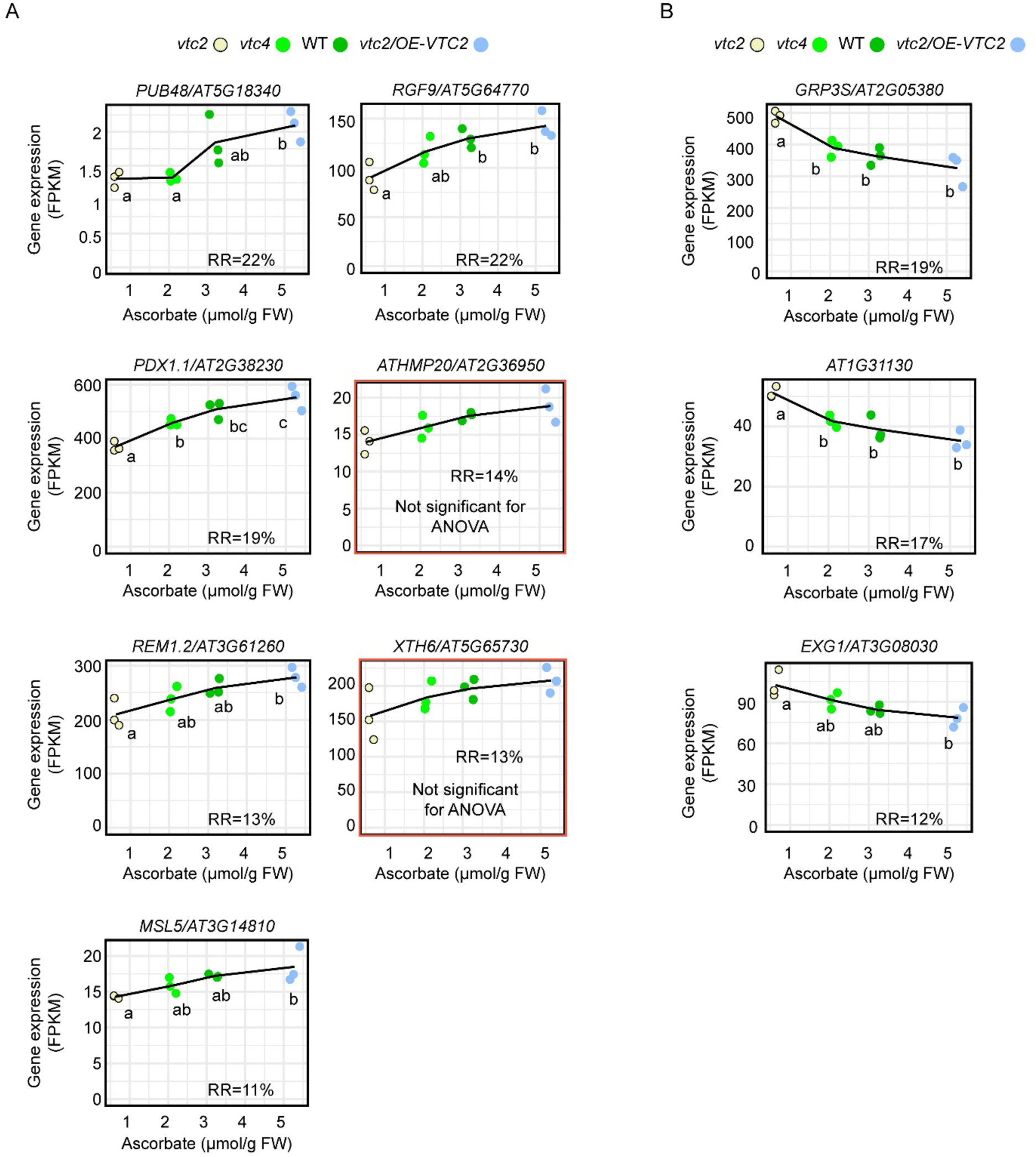
Expression profiles of A) Positively Correlated Genes (PCGs) and B) Negatively Correlated Genes (NCGs) with response to ascorbate (log2Fold-change (FC) *vtc2/OE-VTC2*: log2FC *vtc2*) within the “responding as expected” class, whose sensitivity to ascorbate depletion (FC *vtc4*: FC *vtc2*) falls in “expected” or “high” sensitivity classes. Different letters denote statistically significant differences (One-Way ANOVA, Tukey post-hoc test, α=0.05). Distributions were tested to meet normality (Shapiro–Wilk’s test) and homoscedasticity (Levene’s test) prior to ANOVA. If not meeting any of these requirements, data was transformed using logarithm and, if meeting the requirements, proceeded to perform ANOVA. Otherwise, Kruskal-Wallis followed by Dunn’s post-hoc test were performed. FPKM: Fragments Per Kilobase of transcript per Million mapped reads, FW: fresh weight, RR: relative response as defined in Supp. Fig. 6, *PUB48: PLANT U-BOX 48, RGF9: ROOT MERISTEM GROWTH FACTOR 9, PDX1.1: PYRIDOXINE BIOSYNTHESIS 1.1, XTH6: XYLOGLUCAN ENDOTRANSGLUCOSYLASE/HYDROLASE 6, MSL5: MECHANOSENSITIVE CHANNEL OF SMALL CONDUCTANCE-LIKE 5, ATHMP20: HEAVY METAL ASSOCIATED PROTEIN 20, REM1.2: REMORIN 1.2, AT1G31130: POLYADENYLATE-BINDING PROTEIN 1-B-BINDING PROTEIN, GRP3S: GLYCINE-RICH PROTEIN 3 SHORT ISOFORM, EXG1: ENHANCED XYLEM AND GRAFTING 1*.

**Supplementary Figure S10.**
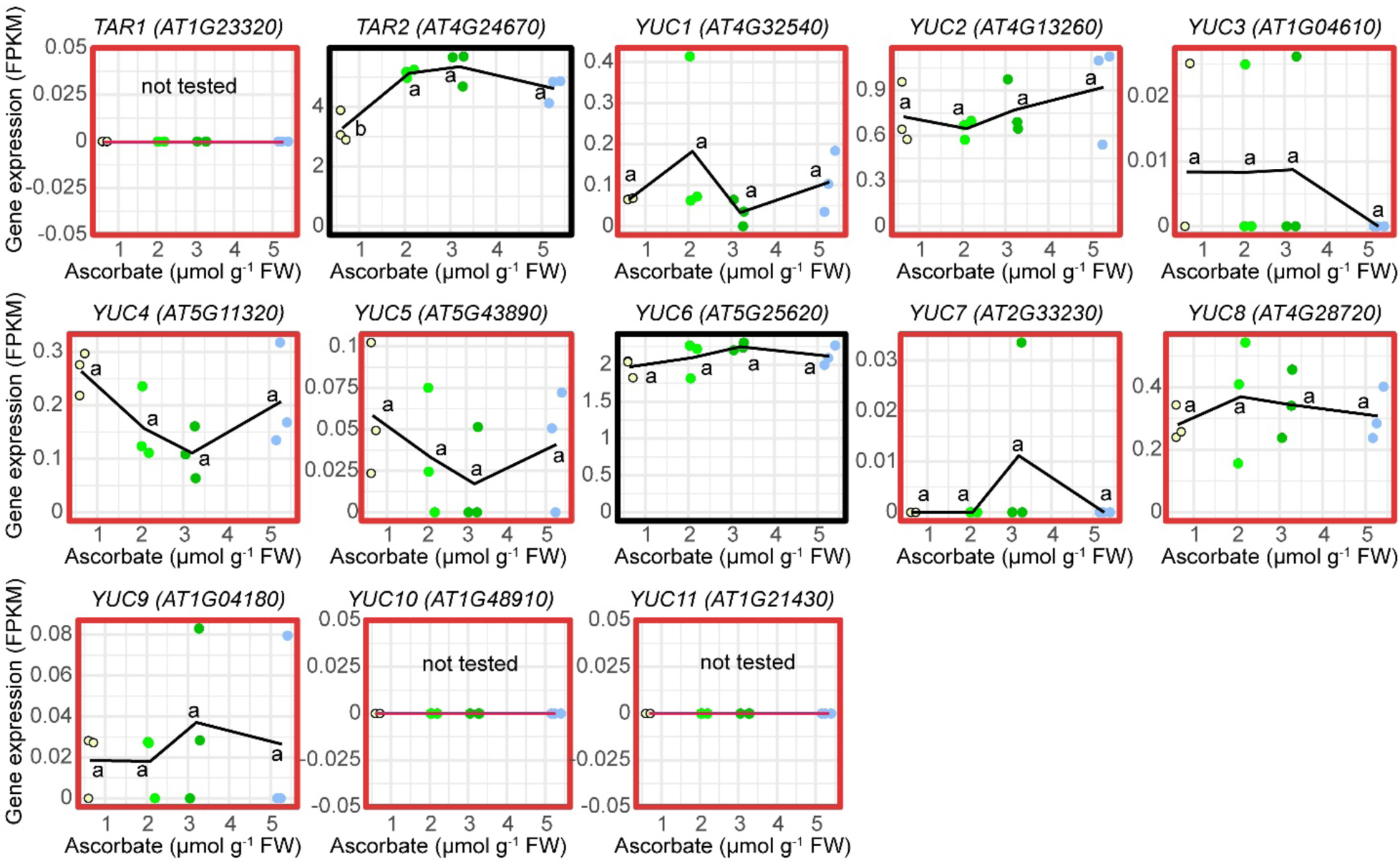
Expression profiles of auxin biosynthesis genes that along with *TAA1* function in the indole pyruvic acid route of IAA production. RNA-seq Fragments Per Kilobase of transcript per Million mapped reads (FPKM) (y-axes) vs ascorbate concentrations (x-axes) are plotted. Plots for genes with very low expression values (FPKM<1) are framed in red and should not be considered reliable, with only *TAR2* and *YUC6* profiles framed in black being potentially informative. Different letters denote statistically significant differences (One-Way ANOVA, Tukey post-hoc test, α=0.05). Distributions were tested to meet normality (Shapiro–Wilk’s test) and homoscedasticity (Levene’s test) prior to ANOVA. If not meeting any of these requirements, data was transformed using logarithm and, if meeting the requirements, proceeded to perform ANOVA. Otherwise, Kruskal-Wallis followed by Dunn’s post-hoc test were performed. FPKM: Fragments Per Kilobase of transcript per Million mapped reads, FW: fresh weight. *TAR1: TAA1-RELATED1, YUC: YUCCA*.

**Supplementary Figure S11.**
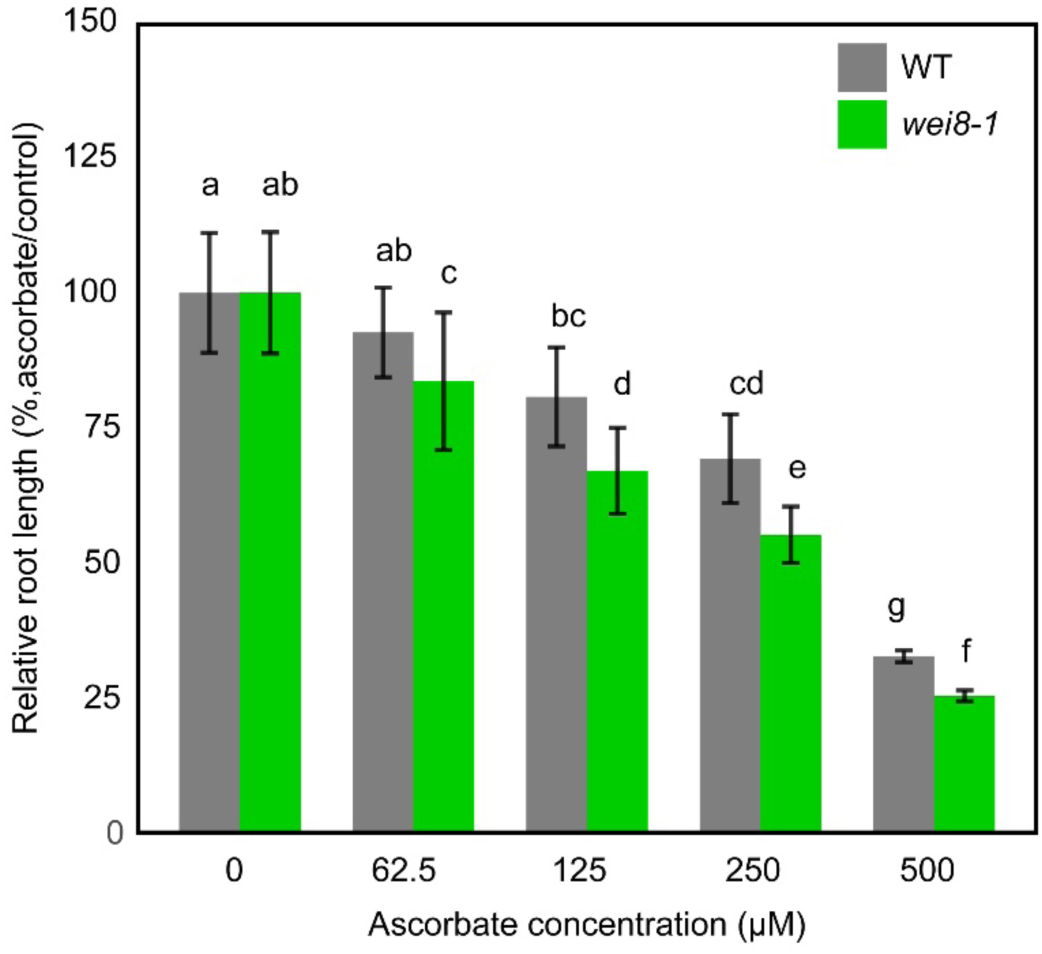
Root sensitivity of mild auxin deficient mutant *wei8* to exogenous ascorbate is higher than WT in five-day-old light-grown seedlings. Roots (n>10) of five-day-old light-grown seedlings on control or ascorbate-supplemented horizontal plates were imaged and length was quantified using FIJI. Different letters denote statistically significant differences (One-Way ANOVA, Tukey post-hoc test, α=0.05). Data was transformed using logarithm to meet homoscedasticity (Levene’s test) and power>0.8 prior to perform ANOVA.

**Supplementary Figure S12.**
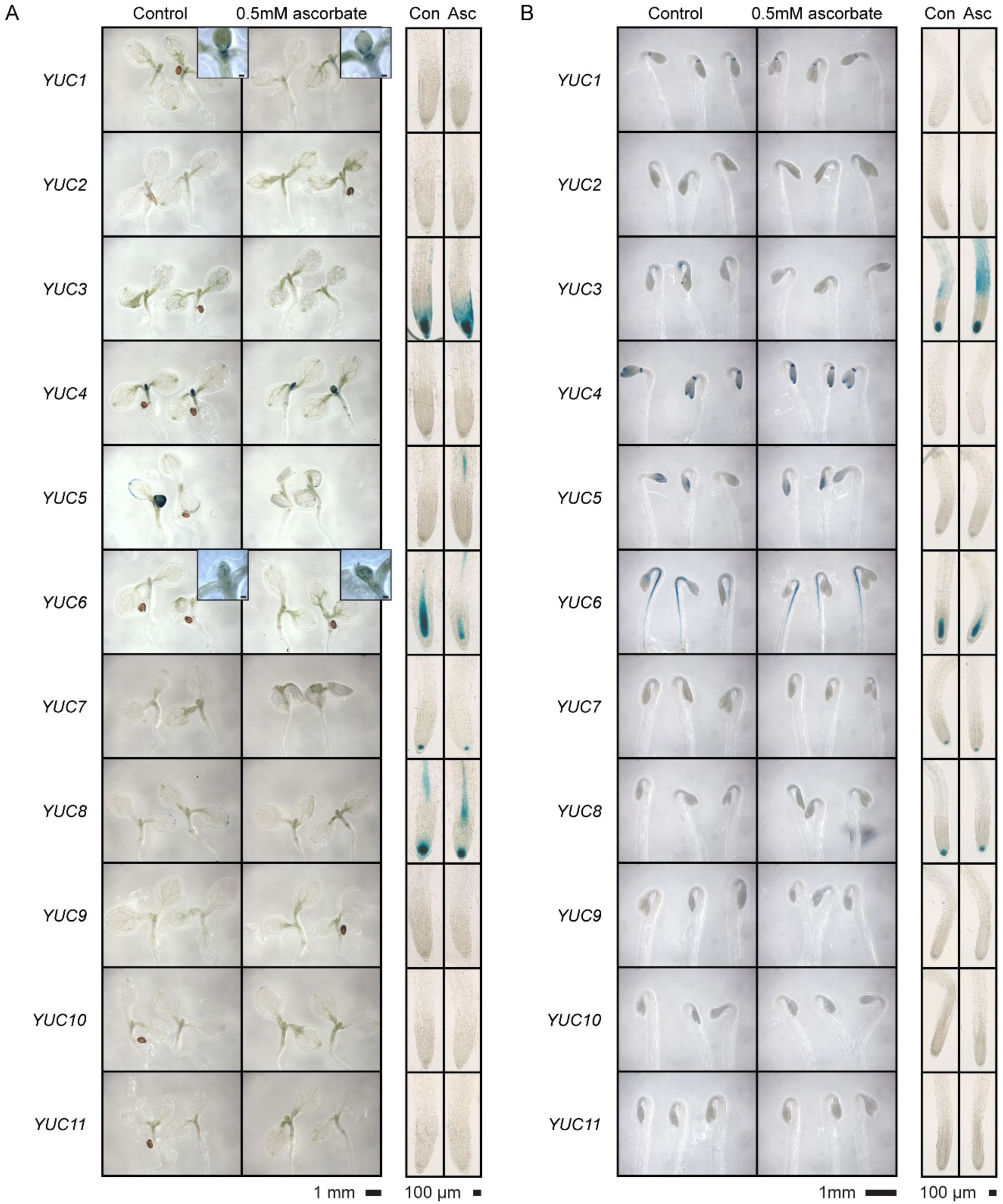
Ascorbate supplementation alters the expression patterns of auxin biosynthesis genes *YUC5*, *YUC6*, and *YUC8* in roots of light-grown seedlings and of *YUC3* in roots of dark-grown seedlings. A,B) Arabidopsis lines expressing recombineering-generated *GUS* translational fusions of *YUC* genes (Brumos et al., 2020) were grown for 5 days under continuous light (A) and for 3 days in the dark (B) and stained for GUS.

**Supplementary Figure S13.**
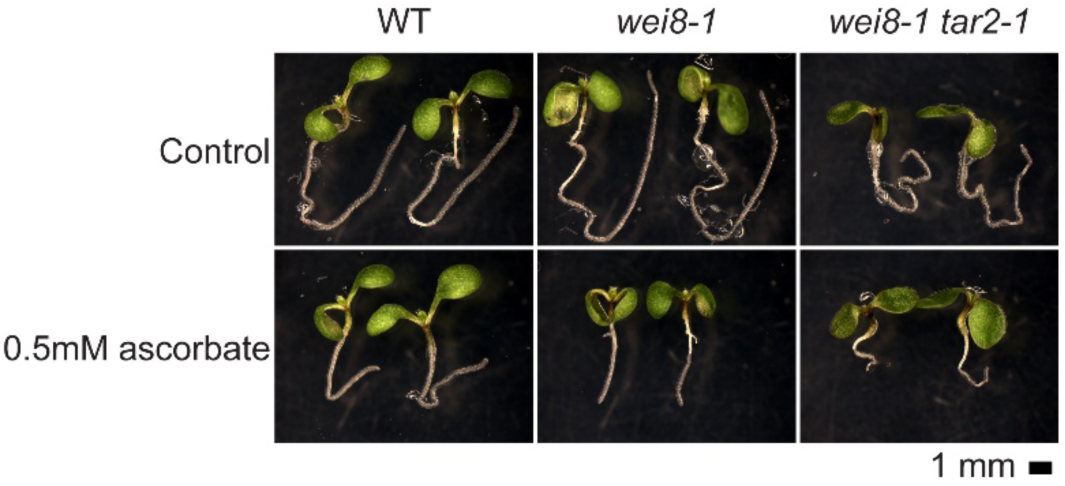
Auxin-deficient double mutant *wei8 tar2* is severely impaired in tryptophan aminotransferase activity, translating into exaggerated shortening of the root in the presence of ascorbate.

**Supplementary Figure S14.**
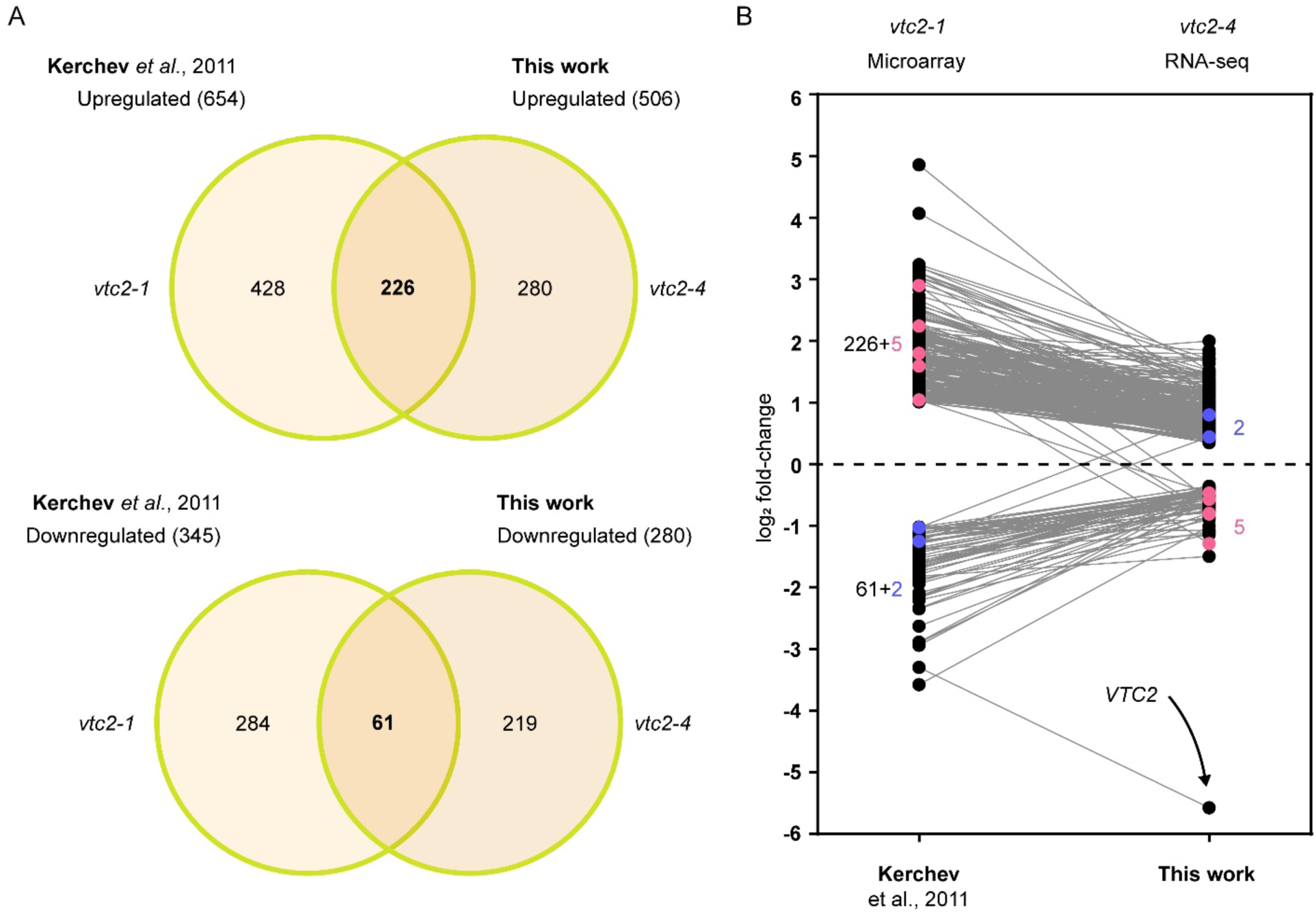
Comparison of *vtc2-*regulated gene lists identified in this work and that published by Kerchev et al. (2011). A) Overlap between upregulated or downregulated genes identified in *vtc2-1* (EMS allele, microarray; Kerchev et al., 2001) and *vtc2-4* (T-DNA allele, RNA-seq; this work). B) Expression fold-change of overlapping genes between the two compared studies (266 upregulated, and 61 downregulated) including DEGs whose fold change is reversed between studies: in pink, 5 genes that were upregulated in Kerchev’s study were downregulated in ours; In blue, 2 genes that were downregulated in Kerchev’s study were upregulated in ours. Data available in Supplementary Material 10.

## References

Aarabi F, Ghigi A, Ahchige MW, Bulut M, Geigenberger P, Neuhaus HE, Sampathkumar A, Alseekh S, Fernie AR (2023) Genome-wide association study unveils ascorbate regulation by PAS/LOV PROTEIN during high light acclimation. Plant Physiol 193: 2037–2054

Ahmad Z, Ramakrishnan M, Wang C, Rehman S, Shahzad A, Wei Q (2024) Unravelling the role of WRKY transcription factors in leaf senescence: Genetic and molecular insights. J Adv Res. doi: 10.1016/j.jare.2024.09.026

Akram NA, Shafiq F, Ashraf M (2017) Ascorbic Acid-A Potential Oxidant Scavenger and Its Role in Plant Development and Abiotic Stress Tolerance. Front Plant Sci 8: 613

Baldet P, Mori K, Decros G, Beauvoit B, Colombié S, Prigent S, Pétriacq P, Gibon Y (2024) Multi-regulated GDP-l-galactose phosphorylase calls the tune in ascorbate biosynthesis. J Exp Bot 75: 2631–2643

Barth C, Moeder W, Klessig DF, Conklin PL (2004) The Timing of Senescence and Response to Pathogens Is Altered in the Ascorbate-Deficient Arabidopsis Mutant vitamin c-1. Plant Physiol 134: 1784–1792

Bengtsson H (2025) R.utils: Various Programming Utilities.

Bennett MJ, Marchant A, Green HG, May ST, Ward SP, Millner PA, Walker AR, Schulz B, Feldmann KA (1996) Arabidopsis AUX1 gene: a permease-like regulator of root gravitropism. Science 273: 948–950

Bian Z, Gao H, Wang C (2020) NAC Transcription Factors as Positive or Negative Regulators during Ongoing Battle between Pathogens and Our Food Crops. Int J Mol Sci 22: 81

Birkenbihl RP, Kracher B, Roccaro M, Somssich IE (2017) Induced Genome-Wide Binding of Three Arabidopsis WRKY Transcription Factors during Early MAMP-Triggered Immunity. Plant Cell 29: 20–38

Blaschke K, Ebata KT, Karimi MM, Zepeda-Martínez JA, Goyal P, Mahapatra S, Tam A, Laird DJ, Hirst M, Rao A, et al (2013) Vitamin C induces Tet-dependent DNA demethylation and a blastocyst-like state in ES cells. Nature 500: 222–226

Bolger AM, Lohse M, Usadel B (2014) Trimmomatic: a flexible trimmer for Illumina sequence data. Bioinforma Oxf Engl 30: 2114–2120

Bournonville C, Mori K, Deslous P, Decros G, Blomeier T, Mauxion J-P, Jorly J, Gadin S, Cassan C, Maucourt M, et al (2023) Blue light promotes ascorbate synthesis by deactivating the PAS/LOV photoreceptor that inhibits GDP-L-galactose phosphorylase. Plant Cell 35: 2615–2634

Brumos J, Robles LM, Yun J, Vu TC, Jackson S, Alonso JM, Stepanova AN (2018) Local Auxin Biosynthesis Is a Key Regulator of Plant Development. Dev Cell 47: 306–318.e5

Brumos J, Zhao C, Gong Y, Soriano D, Patel AP, Perez-Amador MA, Stepanova AN, Alonso JM (2020) An Improved Recombineering Toolset for Plants. Plant Cell 32: 100–122

Bulley SM, Cooney JM, Laing W (2021) Elevating Ascorbate in Arabidopsis Stimulates the Production of Abscisic Acid, Phaseic Acid, and to a Lesser Extent Auxin (IAA) and Jasmonates, Resulting in Increased Expression of DHAR1 and Multiple Transcription Factors Associated with Abiotic Stress Tolerance. Int J Mol Sci 22: 6743

Bulley SM, Rassam M, Hoser D, Otto W, Schünemann N, Wright M, MacRae E, Gleave A, Laing W (2009) Gene expression studies in kiwifruit and gene over-expression in Arabidopsis indicates that GDP-L-galactose guanyltransferase is a major control point of vitamin C biosynthesis. J Exp Bot 60: 765–778

Champely S, Ekstrom C, Dalgaard P, Gill J, Weibelzahl S, Anandkumar A, Ford C, Volcic R, Rosario HD (2020) pwr: Basic Functions for Power Analysis.

Chatterjee IB (1973) Evolution and the biosynthesis of ascorbic acid. Science 182: 1271–1272

Chen M, Dai Y, Liao J, Wu H, Lv Q, Huang Y, Liu L, Feng Y, Lv H, Zhou B, et al (2024) TARGET OF MONOPTEROS: key transcription factors orchestrating plant development and environmental response. J Exp Bot 75: 2214–2234

Chen X, Li C, Wang H, Guo Z (2019) WRKY transcription factors: evolution, binding, and action. Phytopathol Res 1: 13

Clough SJ, Bent AF (1998) Floral dip: a simplified method for -mediated transformation of. Plant J 16: 735–743

Conklin PL, DePaolo D, Wintle B, Schatz C, Buckenmeyer G (2013) Identification of Arabidopsis VTC3 as a putative and unique dual function protein kinase::protein phosphatase involved in the regulation of the ascorbic acid pool in plants. J Exp Bot 64: 2793–2804

Davuluri RV, Sun H, Palaniswamy SK, Matthews N, Molina C, Kurtz M, Grotewold E (2003) AGRIS: Arabidopsis gene regulatory information server, an information resource of Arabidopsis cis-regulatory elements and transcription factors. BMC Bioinformatics 4: 25

Daxinger L, Hunter B, Sheikh M, Jauvion V, Gasciolli V, Vaucheret H, Matzke M, Furner I (2008) Unexpected silencing effects from T-DNA tags in Arabidopsis. Trends Plant Sci 13: 4–6

Delarue M, Prinsen E, Va H, Onckelen, Caboche M, Bellini C (1998) Sur2 mutations of Arabidopsis thaliana define a new locus involved in the control of auxin homeostasis. Plant J 14: 603–611

Dinno A (2024) dunn.test: Dunn’s Test of Multiple Comparisons Using Rank Sums.

Dowdle J, Ishikawa T, Gatzek S, Rolinski S, Smirnoff N (2007) Two genes in Arabidopsis thaliana encoding GDP-L-galactose phosphorylase are required for ascorbate biosynthesis and seedling viability. Plant J Cell Mol Biol 52: 673–689

Edgar R, Domrachev M, Lash AE (2002) Gene Expression Omnibus: NCBI gene expression and hybridization array data repository. Nucleic Acids Res 30: 207–210

Endres S, Tenhaken R (2009) Myoinositol oxygenase controls the level of myoinositol in Arabidopsis, but does not increase ascorbic acid. Plant Physiol 149: 1042–1049

Endrizzi K, Moussian B, Haecker A, Levin JZ, Laux T (1996) The SHOOT MERISTEMLESS gene is required for maintenance of undifferentiated cells in Arabidopsis shoot and floral meristems and acts at a different regulatory level than the meristem genes WUSCHEL and ZWILLE. Plant J Cell Mol Biol 10: 967–979

Fenech M, Amaya I, Valpuesta V, Botella MA (2019) Vitamin C Content in Fruits: Biosynthesis and Regulation. Front Plant Sci. doi: 10.3389/fpls.2018.02006

Fenech M, Amorim-Silva V, Esteban Del Valle A, Arnaud D, Ruiz-Lopez N, Castillo AG, Smirnoff N, Botella MA (2021) The role of GDP-l-galactose phosphorylase in the control of ascorbate biosynthesis. Plant Physiol 185: 1574–1594

Footitt S, Walley PG, Lynn JR, Hambidge AJ, Penfield S, Finch-Savage WE (2020) Trait analysis reveals DOG1 determines initial depth of seed dormancy, but not changes during dormancy cycling that result in seedling emergence timing. New Phytol 225: 2035–2047

Fox J, Weisberg S (2019) An R Companion to Applied Regression, Third. Sage, Thousand Oaks, CA

Foyer CH, Kunert K (2024) The ascorbate–glutathione cycle coming of age. J Exp Bot 75: 2682–2699

Gao Y, Nishikawa H, Badejo AA, Shibata H, Sawa Y, Nakagawa T, Maruta T, Shigeoka S, Smirnoff N, Ishikawa T (2011) Expression of aspartyl protease and C3HC4-type RING zinc finger genes are responsive to ascorbic acid in Arabidopsis thaliana. J Exp Bot 62: 3647–3657

Garnier S, Ross N, Rudis B, Sciaini M, Camargo AP, Scherer C (2024) viridis: Colorblind-Friendly Color Maps for R.

Gautrat P, Buti S, Romanowski A, Lammers M, Matton SEA, Buijs G, Pierik R (2024) Phytochrome-dependent responsiveness to root-derived cytokinins enables coordinated elongation responses to combined light and nitrate cues. Nat Commun 15: 8489

Ge SX, Son EW, Yao R (2018) iDEP: an integrated web application for differential expression and pathway analysis of RNA-Seq data. BMC Bioinformatics 19: 534

Goff L, Trapnell C, Kelley D (2014) cummeRbund: Analysis, exploration, manipulation, and visualization of Cufflinks high-throughput sequencing data.

Graeber K, Linkies A, Steinbrecher T, Mummenhoff K, Tarkowská D, Turečková V, Ignatz M, Sperber K, Voegele A, de Jong H, et al (2014) DELAY OF GERMINATION 1 mediates a conserved coat-dormancy mechanism for the temperature- and gibberellin-dependent control of seed germination. Proc Natl Acad Sci 111: E3571–E3580

Grau J, Franco-Zorrilla JM (2022) TDTHub, a web server tool for the analysis of transcription factor binding sites in plants. Plant J Cell Mol Biol 111: 1203–1215

Graves S, Dorai-Raj H-PP and LS with help from S (2024) multcompView: Visualizations of Paired Comparisons.

Gu Z (2022) Complex heatmap visualization. iMeta 1: e43

Gu Z, Eils R, Schlesner M (2016) Complex heatmaps reveal patterns and correlations in multidimensional genomic data. Bioinformatics 32: 2847–2849

Heberle H, Meirelles GV, da Silva FR, Telles GP, Minghim R (2015) InteractiVenn: a web-based tool for the analysis of sets through Venn diagrams. BMC Bioinformatics 16: 169

Horikoshi M, Tangaut Y, cre Dickey A, Grenié M, Thompson R, Selzer L, Strbenac D, Voronin K, Pulatov D (2024) ggfortify: Data Visualization Tools for Statistical Analysis Results.

Hu L, Lu J, Cheng J, Rao Q, Li Z, Hou H, Lou Z, Zhang L, Li W, Gong W, et al (2015) Structural insight into substrate preference for TET-mediated oxidation. Nature 527: 118–122

Hu Y, Dong Q, Yu D (2012) *Arabidopsis* WRKY46 coordinates with WRKY70 and WRKY53 in basal resistance against pathogen *Pseudomonas syringae*. Plant Sci 185–186: 288–297

Javed T, Gao S-J (2023) WRKY transcription factors in plant defense. Trends Genet 39: 787–801

Jiang K, Meng YL, Feldman LJ (2003) Quiescent center formation in maize roots is associated with an auxin-regulated oxidizing environment. Dev Camb Engl 130: 1429–1438

Kassambara A (2023a) ggpubr: “ggplot2” Based Publication Ready Plots.

Kassambara A (2023b) rstatix: Pipe-Friendly Framework for Basic Statistical Tests.

Kawade K, Horiguchi G, Tsukaya H (2010) Non-cell-autonomously coordinated organ size regulation in leaf development. Dev Camb Engl 137: 4221–4227

Kerchev PI, Pellny TK, Vivancos PD, Kiddle G, Hedden P, Driscoll S, Vanacker H, Verrier P, Hancock RD, Foyer CH (2011) The transcription factor ABI4 Is required for the ascorbic acid-dependent regulation of growth and regulation of jasmonate-dependent defense signaling pathways in Arabidopsis. Plant Cell 23: 3319–3334

Kerk NM, Jiang K, Feldman LJ (2000) Auxin Metabolism in the Root Apical Meristem. Plant Physiol 122: 925–932

Kim BC, Soh MS, Kang BJ, Furuya M, Nam HG (1996) Two dominant photomorphogenic mutations of Arabidopsis thaliana identified as suppressor mutations of hy2. Plant J 9: 441–456

Kim D, Langmead B, Salzberg SL (2015) HISAT: a fast spliced aligner with low memory requirements. Nat Methods 12: 357–360

Korbei B, Moulinier-Anzola J, De-Araujo L, Lucyshyn D, Retzer K, Khan MA, Luschnig C (2013) *Arabidopsis* TOL Proteins Act as Gatekeepers for Vacuolar Sorting of PIN2 Plasma Membrane Protein. Curr Biol 23: 2500–2505

Krüger T, Brandt D, Sodenkamp J, Gasper M, Romera-Branchat M, Ahloumessou F, Gehring E, Drotleff J, Bell C, Kramer K, et al (2025) DOG1 controls dormancy independently of ABA core signaling kinases regulation by preventing AFP dephosphorylation through AHG1. Sci Adv 11: eadr8502

Kubalová M, Müller K, Dobrev PI, Rizza A, Jones AM, Fendrych M (2024) Auxin co-receptor IAA17/AXR3 controls cell elongation in Arabidopsis thaliana root solely by modulation of nuclear auxin pathway. New Phytol 241: 2448–2463

Laing WA, Martínez-Sánchez M, Wright MA, Bulley SM, Brewster D, Dare AP, Rassam M, Wang D, Storey R, Macknight RC, et al (2015) An upstream open reading frame is essential for feedback regulation of ascorbate biosynthesis in Arabidopsis. Plant Cell 27: 772–786

Lee Y, Kim MW, Kim SH (2007) Cell type identity inArabidopsis roots is altered by both ascorbic acid-induced changes in the redox environment and the resultant endogenous auxin response. J Plant Biol 50: 484–489

Lee Y, Park CH, Ram Kim A, Chang SC, Kim S-H, Lee WS, Kim S-K (2011) The effect of ascorbic acid and dehydroascorbic acid on the root gravitropic response in Arabidopsis thaliana. Plant Physiol Biochem PPB 49: 909–916

Leyser HM, Pickett FB, Dharmasiri S, Estelle M (1996) Mutations in the AXR3 gene of Arabidopsis result in altered auxin response including ectopic expression from the SAUR-AC1 promoter. Plant J Cell Mol Biol 10: 403–413

Lim B, Smirnoff N, Cobbett CS, Golz JF (2016) Ascorbate-Deficient vtc2 Mutants in Arabidopsis Do Not Exhibit Decreased Growth. Front Plant Sci 7: 1025

Linster CL, Gomez TA, Christensen KC, Adler LN, Young BD, Brenner C, Clarke SG (2007) Arabidopsis VTC2 encodes a GDP-L-galactose phosphorylase, the last unknown enzyme in the Smirnoff-Wheeler pathway to ascorbic acid in plants. J Biol Chem 282: 18879–18885

Lipner S (2018) A classic case of scurvy. Lancet Lond Engl 392: 431

Liu S, Kracher B, Ziegler J, Birkenbihl RP, Somssich IE (2015) Negative regulation of ABA signaling by WRKY33 is critical for Arabidopsis immunity towards Botrytis cinerea 2100. eLife 4: e07295

Lorence A, Chevone BI, Mendes P, Nessler CL (2004) myo-inositol oxygenase offers a possible entry point into plant ascorbate biosynthesis. Plant Physiol 134: 1200–1205

Lukowitz W, Nickle TC, Meinke DW, Last RL, Conklin PL, Somerville CR (2001) Arabidopsis cyt1 mutants are deficient in a mannose-1-phosphate guanylyltransferase and point to a requirement of N-linked glycosylation for cellulose biosynthesis. Proc Natl Acad Sci U S A 98: 2262–2267

Mangano S, Denita-Juarez SP, Choi H-S, Marzol E, Hwang Y, Ranocha P, Velasquez SM, Borassi C, Barberini ML, Aptekmann AA, et al (2017) Molecular link between auxin and ROS-mediated polar growth. Proc Natl Acad Sci U S A 114: 5289–5294

Marchant A, Bhalerao R, Casimiro I, Eklöf J, Casero PJ, Bennett M, Sandberg G (2002) AUX1 promotes lateral root formation by facilitating indole-3-acetic acid distribution between sink and source tissues in the Arabidopsis seedling. Plant Cell 14: 589–597

Marchant A, Kargul J, May ST, Muller P, Delbarre A, Perrot-Rechenmann C, Bennett MJ (1999) AUX1 regulates root gravitropism in Arabidopsis by facilitating auxin uptake within root apical tissues. EMBO J 18: 2066–2073

Mi H, Muruganujan A, Huang X, Ebert D, Mills C, Guo X, Thomas PD (2019) Protocol Update for large-scale genome and gene function analysis with the PANTHER classification system (v.14.0). Nat Protoc 14: 703–721

Milne I, Stephen G, Bayer M, Cock PJA, Pritchard L, Cardle L, Shaw PD, Marshall D (2013) Using Tablet for visual exploration of second-generation sequencing data. Brief Bioinform 14: 193–202

Minor EA, Court BL, Young JI, Wang G (2013) Ascorbate induces ten-eleven translocation (Tet) methylcytosine dioxygenase-mediated generation of 5-hydroxymethylcytosine. J Biol Chem 288: 13669–13674

Mlotshwa S, Pruss GJ, Gao Z, Mgutshini NL, Li J, Chen X, Bowman LH, Vance V (2010) Transcriptional Silencing Induced by Arabidopsis T-DNA Mutants is Associated with 35S Promoter siRNAs and Requires Genes Involved in siRNA-mediated Chromatin Silencing. Plant J Cell Mol Biol 64: 699–704

Mounet-Gilbert L, Dumont M, Ferrand C, Bournonville C, Monier A, Jorly J, Lemaire-Chamley M, Mori K, Atienza I, Hernould M, et al (2016) Two tomato GDP-D-mannose epimerase isoforms involved in ascorbate biosynthesis play specific roles in cell wall biosynthesis and development. J Exp Bot 67: 4767–4777

Mukherjee M, Larrimore KE, Ahmed NJ, Bedick TS, Barghouthi NT, Traw MB, Barth C (2010) Ascorbic acid deficiency in arabidopsis induces constitutive priming that is dependent on hydrogen peroxide, salicylic acid, and the NPR1 gene. Mol Plant-Microbe Interact MPMI 23: 340– 351

Nakagawa T, Kurose T, Hino T, Tanaka K, Kawamukai M, Niwa Y, Toyooka K, Matsuoka K, Jinbo T, Kimura T (2007) Development of series of gateway binary vectors, pGWBs, for realizing efficient construction of fusion genes for plant transformation. J Biosci Bioeng 104: 34–41

Neuteboom LW, Ng JM, Kuyper M, Clijdesdale OR, Hooykaas PJ, van der Zaal BJ (1999) Isolation and characterization of cDNA clones corresponding with mRNAs that accumulate during auxin-induced lateral root formation. Plant Mol Biol 39: 273–287

Nguyen NH, Sng BJR, Chin HJ, Choi IKY, Yeo HC, Jang I-C (2023) HISTONE DEACETYLASE 9 promotes hypocotyl-specific auxin response under shade. Plant J 116: 804–822

Norris SR, Shen X, Della Penna D (1998) Complementation of the Arabidopsis pds1 Mutation with the Gene Encoding p-Hydroxyphenylpyruvate Dioxygenase. Plant Physiol 117: 1317–1323

O’Malley RC, Huang S-SC, Song L, Lewsey MG, Bartlett A, Nery JR, Galli M, Gallavotti A, Ecker JR (2016) Cistrome and Epicistrome Features Shape the Regulatory DNA Landscape. Cell 165: 1280–1292

Padayatty SJ, Levine M (2016) Vitamin C: the known and the unknown and Goldilocks. Oral Dis 22: 463–493

Page M, Sultana N, Paszkiewicz K, Florance H, Smirnoff N (2012) The influence of ascorbate on anthocyanin accumulation during high light acclimation in Arabidopsis thaliana: further evidence for redox control of anthocyanin synthesis. Plant Cell Environ 35: 388–404

Palaniswamy SK, James S, Sun H, Lamb RS, Davuluri RV, Grotewold E (2006) AGRIS and AtRegNet. a platform to link cis-regulatory elements and transcription factors into regulatory networks. Plant Physiol 140: 818–829

Pastor V, Luna E, Ton J, Cerezo M, García-Agustín P, Flors V (2013) Fine tuning of reactive oxygen species homeostasis regulates primed immune responses in Arabidopsis. Mol Plant-Microbe Interact MPMI 26: 1334–1344

Pastori GM, Kiddle G, Antoniw J, Bernard S, Veljovic-Jovanovic S, Verrier PJ, Noctor G, Foyer CH (2003) Leaf Vitamin C Contents Modulate Plant Defense Transcripts and Regulate Genes That Control Development through Hormone Signaling. Plant Cell 15: 939–951

Pavet V, Olmos E, Kiddle G, Mowla S, Kumar S, Antoniw J, Alvarez ME, Foyer CH (2005) Ascorbic acid deficiency activates cell death and disease resistance responses in Arabidopsis. Plant Physiol 139: 1291–1303

Pineau B, Layoune O, Danon A, De Paepe R (2008) l-Galactono-1,4-lactone Dehydrogenase Is Required for the Accumulation of Plant Respiratory Complex I*. J Biol Chem 283: 32500–32505

Qi T, Liu Z, Fan M, Chen Y, Tian H, Wu D, Gao H, Ren C, Song S, Xie D (2017) GDP-D-mannose epimerase regulates male gametophyte development, plant growth and leaf senescence in Arabidopsis. Sci Rep 7: 10309

R Core Team (2025) R: A Language and Environment for Statistical Computing.

Ran X, Zhao F, Wang Y, Liu J, Zhuang Y, Ye L, Qi M, Cheng J, Zhang Y (2020) Plant Regulomics: a data-driven interface for retrieving upstream regulators from plant multi-omics data. Plant J 101: 237–248

Reikersdorfer KN, Singh A, Young JD, Batty MB, Steele AE, Yuen LC, Momtaz DA, Weissert JN, Liu DS, Hogue GD (2024) The Troubling Rise of Scurvy: A Review and National Analysis of Incidence, Associated Risk Factors, and Clinical Manifestations. J Am Acad Orthop Surg Glob Res Rev 8: e24.00162

Reuber TL, Ausubel FM (1996) Isolation of Arabidopsis genes that differentiate between resistance responses mediated by the RPS2 and RPM1 disease resistance genes. Plant Cell 8: 241–249

Romano CP, Robson PR, Smith H, Estelle M, Klee H (1995) Transgene-mediated auxin overproduction in Arabidopsis: hypocotyl elongation phenotype and interactions with the hy6-1 hypocotyl elongation and axr1 auxin-resistant mutants. Plant Mol Biol 27: 1071–1083

Seo PJ, Lee A-K, Xiang F, Park C-M (2008) Molecular and functional profiling of Arabidopsis pathogenesis-related genes: insights into their roles in salt response of seed germination. Plant Cell Physiol 49: 334–344

Smirnoff N (2018) Ascorbic acid metabolism and functions: A comparison of plants and mammals. Free Radic Biol Med 122: 116–129

Smirnoff N, Wheeler GL (2024) The ascorbate biosynthesis pathway in plants is known, but there is a way to go with understanding control and functions. J Exp Bot 75: 2604–2630

Spurney RJ, Van den Broeck L, Clark NM, Fisher AP, de Luis Balaguer MA, Sozzani R (2020) tuxnet: a simple interface to process RNA sequencing data and infer gene regulatory networks. Plant J 101: 716–730

Stepanova AN, Hoyt JM, Hamilton AA, Alonso JM (2005) A Link between Ethylene and Auxin Uncovered by the Characterization of Two Root-Specific Ethylene-Insensitive Mutants in Arabidopsis. Plant Cell 17: 2230–2242

Stepanova AN, Robertson-Hoyt J, Yun J, Benavente LM, Xie D-Y, Doležal K, Schlereth A, Jürgens G, Alonso JM (2008) TAA1-Mediated Auxin Biosynthesis Is Essential for Hormone Crosstalk and Plant Development. Cell 133: 177–191

Stepanova AN, Yun J, Robles LM, Novak O, He W, Guo H, Ljung K, Alonso JM (2011) The *Arabidopsis* YUCCA1 Flavin Monooxygenase Functions in the Indole-3-Pyruvic Acid Branch of Auxin Biosynthesis. Plant Cell 23: 3961–3973

Swarup R, Bhosale R (2019) Developmental Roles of AUX1/LAX Auxin Influx Carriers in Plants. Front Plant Sci 10: 1306

Szecowka M, Heise R, Tohge T, Nunes-Nesi A, Vosloh D, Huege J, Feil R, Lunn J, Nikoloski Z, Stitt M, et al (2013) Metabolic Fluxes in an Illuminated Arabidopsis Rosette. Plant Cell 25: 694–714

Tao Y, Ferrer J-L, Ljung K, Pojer F, Hong F, Long JA, Li L, Moreno JE, Bowman ME, Ivans LJ, et al (2008) Rapid Synthesis of Auxin via a New Tryptophan-Dependent Pathway Is Required for Shade Avoidance in Plants. Cell 133: 164–176

Thomas PD, Ebert D, Muruganujan A, Mushayahama T, Albou L-P, Mi H (2022) PANTHER: Making genome-scale phylogenetics accessible to all. Protein Sci Publ Protein Soc 31: 8–22

Thomma BP, Eggermont K, Penninckx IA, Mauch-Mani B, Vogelsang R, Cammue BP, Broekaert WF (1998) Separate jasmonate-dependent and salicylate-dependent defense-response pathways in Arabidopsis are essential for resistance to distinct microbial pathogens. Proc Natl Acad Sci U S A 95: 15107–15111

Torabinejad J, Donahue JL, Gunesekera BN, Allen-Daniels MJ, Gillaspy GE (2009) VTC4 is a bifunctional enzyme that affects myoinositol and ascorbate biosynthesis in plants. Plant Physiol 150: 951–961

Tóth SZ, Schansker G, Garab G (2013) The physiological roles and metabolism of ascorbate in chloroplasts. Physiol Plant 148: 161–175

Trapnell C, Hendrickson DG, Sauvageau M, Goff L, Rinn JL, Pachter L (2013) Differential analysis of gene regulation at transcript resolution with RNA-seq. Nat Biotechnol 31: 46–53

Trapnell C, Roberts A, Goff L, Pertea G, Kim D, Kelley DR, Pimentel H, Salzberg SL, Rinn JL, Pachter L (2012) Differential gene and transcript expression analysis of RNA-seq experiments with TopHat and Cufflinks. Nat Protoc 7: 562–578

Tyburski J, Dunajska-Ordak K, Skorupa M, Tretyn A (2012) Role of Ascorbate in the Regulation of the Arabidopsis thaliana Root Growth by Phosphate Availability. J Bot 2012: 580342

Tyystjärvi E (2008) Photoinhibition of Photosystem II and photodamage of the oxygen evolving manganese cluster. Coord Chem Rev 252: 361–376

Uknes S, Mauch-Mani B, Moyer M, Potter S, Williams S, Dincher S, Chandler D, Slusarenko A, Ward E, Ryals J (1992) Acquired resistance in Arabidopsis. Plant Cell 4: 645–656

Ulmasov T, Murfett J, Hagen G, Guilfoyle TJ (1997) Aux/IAA proteins repress expression of reporter genes containing natural and highly active synthetic auxin response elements. Plant Cell 9: 1963–1971

Urueña-Palacio S, Ferreyro BL, Fernández-Otero LG, Calo PD (2018) Adult scurvy associated with psychiatric disorders and breast feeding. BMJ Case Rep 2018: bcr2017223686, bcr-2017– 223686

Veljović-Jovanović S, Vidović M, Morina F (2017) Ascorbate as a Key Player in Plant Abiotic Stress Response and Tolerance. In MA Hossain, S Munné-Bosch, DJ Burritt, P Diaz-Vivancos, M Fujita, A Lorence, eds, Ascorbic Acid Plant Growth Dev. Stress Toler. Springer International Publishing, Cham, pp 47–109

Wang Q, Shi H, Huang R, Ye R, Luo Y, Guo Z, Lu S (2021) AIR12 confers cold tolerance through regulation of the CBF cold response pathway and ascorbate homeostasis. Plant Cell Environ 44: 1522–1533

Wang Y, Li Y, Rosas-Diaz T, Caceres-Moreno C, Lozano-Duran R, Macho AP (2019) The IMMUNE-ASSOCIATED NUCLEOTIDE-BINDING 9 Protein Is a Regulator of Basal Immunity in Arabidopsis thaliana. Mol Plant-Microbe Interact MPMI 32: 65–75

Weijers D, Schlereth A, Ehrismann JS, Schwank G, Kientz M, Jürgens G (2006) Auxin triggers transient local signaling for cell specification in Arabidopsis embryogenesis. Dev Cell 10: 265–270

Whitford R, Fernandez A, Tejos R, Pérez AC, Kleine-Vehn J, Vanneste S, Drozdzecki A, Leitner J, Abas L, Aerts M, et al (2012) GOLVEN Secretory Peptides Regulate Auxin Carrier Turnover during Plant Gravitropic Responses. Dev Cell 22: 678–685

Wickham H, Chang W, Henry L, Pedersen TL, Takahashi K, Wilke C, Woo K, Yutani H, Dunnington D, van den Brand T, et al (2025a) ggplot2: Create Elegant Data Visualisations Using the Grammar of Graphics.

Wickham H, François R, Henry L, Müller K, Vaughan D, Software P, PBC (2023) dplyr: A Grammar of Data Manipulation.

Wickham H, Henry L, Pedersen TL, Luciani TJ, Decorde M, Lise V, code) TP (Early line dashing, code) DG (Line dashing code and early raster, implementation) YQ (Improved styles; polypath, code) HM (Opacity, et al (2025b) svglite: An “SVG” Graphics Device.

Wu S-Y, Hou L-L, Zhu J, Wang Y-C, Zheng Y-L, Hou J-Q, Yang Z-N, Lou Y (2023) Ascorbic acid-mediated reactive oxygen species homeostasis modulates the switch from tapetal cell division to cell differentiation in Arabidopsis. Plant Cell 35: 1474–1495

Yamada M, Greenham K, Prigge MJ, Jensen PJ, Estelle M (2009) The TRANSPORT INHIBITOR RESPONSE2 Gene Is Required for Auxin Synthesis and Diverse Aspects of Plant Development. Plant Physiol 151: 168–179

Yilmaz A, Mejia-Guerra MK, Kurz K, Liang X, Welch L, Grotewold E (2011) AGRIS: the Arabidopsis Gene Regulatory Information Server, an update. Nucleic Acids Res 39: D1118–1122

Young JI, Züchner S, Wang G (2015) Regulation of the Epigenome by Vitamin C. Annu Rev Nutr 35: 545–564

Yu Y, Wang J, Li S, Kakan X, Zhou Y, Miao Y, Wang F, Qin H, Huang R (2019) Ascorbic Acid Integrates the Antagonistic Modulation of Ethylene and Abscisic Acid in the Accumulation of Reactive Oxygen Species. Plant Physiol 179: 1861–1875

Yuan X, Wang H, Cai J, Li D, Song F (2019) NAC transcription factors in plant immunity. Phytopathol Res 1: 3

Zhao Y, Christensen SK, Fankhauser C, Cashman JR, Cohen JD, Weigel D, Chory J (2001) A role for flavin monooxygenase-like enzymes in auxin biosynthesis. Science 291: 306–309

Zhao Y, Hull AK, Gupta NR, Goss KA, Alonso J, Ecker JR, Normanly J, Chory J, Celenza JL (2002) Trp-dependent auxin biosynthesis in Arabidopsis: involvement of cytochrome P450s CYP79B2 and CYP79B3. Genes Dev 16: 3100–3112

Zhou Z-Y, Zhang C-G, Wu L, Zhang C-G, Chai J, Wang M, Jha A, Jia P-F, Cui S-J, Yang M, et al (2011) Functional characterization of the CKRC1/TAA1 gene and dissection of hormonal actions in the Arabidopsis root. Plant J Cell Mol Biol 66: 516–527

